# Bright Days Buffer Nighttime Light: Daytime Illumination Shapes Sex Differences in Sleep and Circadian Regulation

**DOI:** 10.64898/2026.02.25.707542

**Authors:** Yumeng Wang, Caihan Tony Chen, Tom Deboer, Gene D. Block, Ketema N. Paul, Christopher S. Colwell

**Affiliations:** Department of Psychiatry & Biobehavioral Sciences, University of California Los Angeles, Los Angeles, CA, USA; Laboratory for Neurophysiology, Department of Cell and Chemical Biology, Leiden University Medical Center, Leiden, The Netherlands; Department of Integrative Biology and Physiology, University of California Los Angeles, Los Angeles, CA, USA

**Keywords:** Sleep, sleep architecture, sex difference, light at night, light intensity, bright light, slow wave activity, circadian disruption

## Abstract

Sex differences in sleep and wakefulness are well documented in humans but remain inconsistent in rodent studies, suggesting strong sensitivity to experimental context. In prior work, we observed no sex differences in sleep-wake architecture under relatively bright daytime light, raising the possibility that daytime illumination is a critical but underappreciated variable shaping sex-dependent sleep regulation. Here, we tested the hypothesis that daytime light intensity modulates sex differences in sleep-wake architecture and vulnerability to dim light at night (DLaN). Male and female C57BL/6J mice were exposed to acute (one night) or chronic (two weeks) DLaN (10 lux) under three daytime light intensities (50, 100, 300 lux). Sleep was assessed using electroencephalographic-based measures of vigilance states and slow wave activity (SWA). Dim daytime light (50 lux) unmasked robust sex differences in dark-phase sleep-wake architecture that were absent under brighter daytime light (300 lux). Acute DLaN reduced early-night wakefulness in both sexes under low daytime light but had minimal effect under bright daytime conditions.

Following chronic DLaN, males exhibited reduced dim light-phase wakefulness and dampened rhythm amplitude, whereas females showed pronounced phase shifts, rhythm attenuation, and altered timing of SWA under 50 and 100 lux. These changes were largely prevented under bright daytime light. Together, these findings identify daytime light intensity as a critical contextual factor governing sex-specific regulation of sleep and vulnerability to nighttime light, providing a unifying framework to reconcile inconsistencies in the rodent sleep literature.

**Highlights:**

- Daytime light intensity shapes sex differences in sleep–wake architecture
- Acute and chronic nighttime light elicit distinct sex-specific sleep responses
- Females exhibit greater circadian and slow-wave vulnerability to nighttime light
- Brighter daytime light buffers sleep and circadian disruption

## Introduction

Sex differences in sleep are robustly documented in humans but remain surprisingly unresolved in animal models. In humans, women more frequently report insomnia symptoms, poorer subjective sleep quality, and greater difficulty maintaining sleep, ^1^ yet objective measures show that women obtain longer total sleep time on average than men.^2^ These convergent findings suggest a strong influence of biological sex on sleep regulation in humans. In rodents, however, the evidence is inconsistent, even within the same mouse strain.^3,4^ Some studies report that males spend less time awake and more time asleep than females, ^3,5–7^ whereas others detect no sex differences in sleep or wakefulness.^4,8^ The source of this variability remains unclear. Notably, most studies do not report or systematically control the daytime lighting environment, despite the central role of light in shaping circadian and sleep–wake regulation.

Our previous work provides a potential clue: under relatively bright daytime light (300 lux), male and female C57BL/6J mice showed no detectable sex differences in sleep architecture.^4^ Bright daytime light strengthens circadian rhythms in human and other species,^9,10^ and because sleep–wake organization is strongly governed by the circadian system,^11^ these findings raise the possibility that daytime light intensity itself may mask or reveal sex differences in sleep. To date, however, this hypothesis has not been directly tested.

Light exposure at night further complicates this picture. Dim light at night (DLaN) is increasingly recognized as a risk factor for sleep disruption, circadian misalignment, cardiometabolic dysfunction, and mood disorders.^12–15^ Human studies suggest that a “bright day & dark night” lighting profile is associated with improved sleep and health outcomes.^9^ Yet rodent studies of DLaN report highly variable outcomes, with some demonstrating pronounced sleep and circadian disruption and others finding minimal effects. A striking but underappreciated feature of this literature is the wide range of daytime light intensities used across studies.

Here, we directly test whether daytime light intensity shapes sex differences in sleep and vulnerability to nighttime light exposure. By systematically manipulating daytime illumination and nighttime light exposure in male and female mice, we test two related hypotheses: first, that sex differences in sleep are contingent on daytime light intensity; and second, that brighter daytime light buffers the effects of nighttime light on sleep and circadian regulation. Together, this work provides a unified framework for reconciling inconsistencies in literature and offers new insight into how lighting environments shape sleep health.

## Result

Reports of sex differences in rodent sleep phenotypes have been inconsistent across studies. In our previous work, we observed no sex differences in sleep–wake rhythms in C57BL/6J mice housed under relatively bright daytime light (300 lux).^4^ Because daytime light intensity is rarely reported and varies substantially across laboratories, we hypothesized that differences in daytime illumination may account for this variability. We therefore examined sleep under three daytime light intensities commonly used in rodent studies: dim (50 lux), moderate (100 lux) and bright (300 lux) to determine whether daytime light modulates sex differences in sleep.

### Daytime light intensity alters sleep quantity but not sleep quality in male and female mice

Daytime light intensity did not significantly affect sleep or wakefulness during the light phase across any lighting condition (**Fig. 1**). In contrast, daytime light intensity strongly influenced the amount of sleep and wakefulness during the dark phase, where sex differences were most pronounced under dim daytime light and progressively diminished with increasing daytime illumination. Across the 24-hour cycle, sex differences were primarily restricted to the dark phase, with no significant differences detected during the light phase. Under 50 lux daytime light condition, female mice exhibited significantly more wakefulness and less sleep during the dark period compared with males, particularly around zeitgeber time (ZT) 18–20 for wakefulness and ZT19–20 for non-rapid eye moment (NREM) sleep. A significant interaction between ZT and sex was observed for wakefulness and NREM sleep, with sex differences evident across all vigilance states (**Fig. 1A–C; Table S1**).

**Figure 1.**
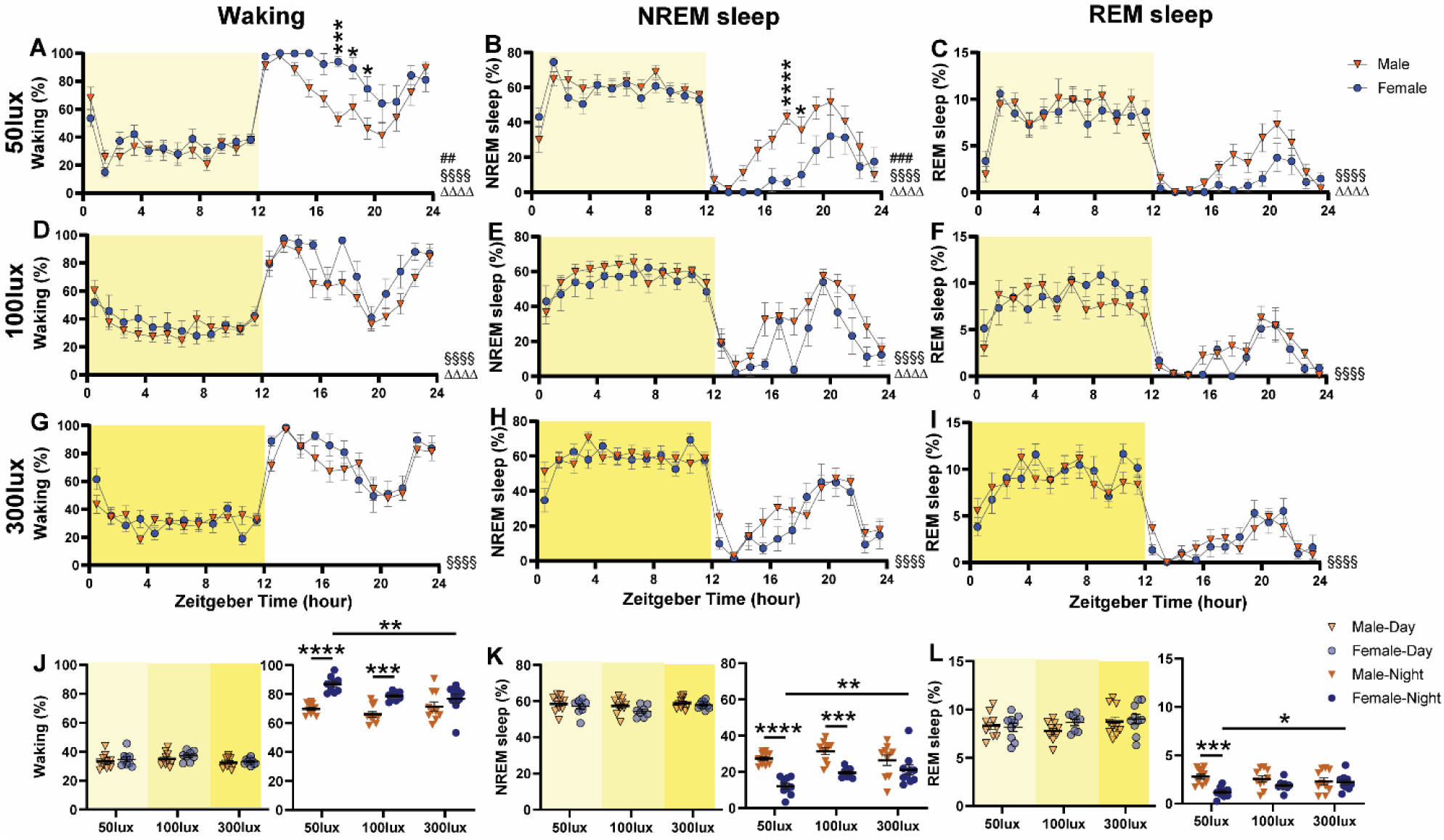
Male and female WT mice exhibit distinct sleep architecture under different daytime light intensities. (**A–I**) 24 hour distribution of time spent in waking (**A, D, G**), NREM sleep (**B, E, H**), and REM sleep (**C, F, I**) in male (orange triangles) and female (blue circles) mice under 50 lux (**A–C**), 100 lux (**D–F**), and 300 lux (**G–I**) daylight conditions. Pound signs (#) indicate a significant interaction between the factors zeitgeber time and sex (two-way repeated-measures ANOVA with Geisser–Greenhouse correction; ## *p* < 0.01, ### *p* < 0.001). Main effects of zeitgeber time and sex are indicated by § and Δ, respectively (§§§§ *p* < 0.0001; ΔΔΔΔ *p* < 0.0001). Asterisks indicate significant post hoc differences between male and female mice (Bonferroni multiple comparisons test; * *p* < 0.05, *** *p* < 0.001, **** *p* < 0.0001). (**J–L**) Daytime (left) and nighttime (right) distribution of time spent in waking, NREM sleep, and REM sleep. Asterisks indicate significant post hoc comparisons (Bonferroni multiple comparisons test; * *p* < 0.05, ** *p* < 0.01, *** *p* < 0.001, **** *p* < 0.0001). Different shades of yellow represent different daytime light intensities. Sample sizes: 50 lux, 10 males and 9 females; 100 lux, 10 males and 8 females; 300 lux, 10 males and 10 females. Data are presented as mean ± SEM.

Under 100 lux and 300 lux daytime light conditions, hourly sleep–wake distributions did not differ significantly between sexes (**Fig. 1D–I**); however, sex differences in overall dark-phase wakefulness and NREM sleep remained detectable under 100 lux light condition (**Fig. 1D, E; Table S1**). In contrast, under bright daytime light (300 lux), only a main effect of ZT was observed, with no significant sex effect or interaction, consistent with our previous findings (**Fig. 1G–I; Table S1**; Wang et al., 2025). Quantification of sleep–wake percentages across light and dark phases confirmed these findings. Under 50 lux daytime light, female mice spent significantly more time awake and less time in NREM and rapid eye moment (REM) sleep during the dark phase compared with males (**Fig. 1J–L**). Although hourly differences were not detected under 100 lux, females still exhibited increased dark-phase wakefulness and reduced NREM sleep relative to males. No sex differences were observed under 300 lux daytime light (**Fig. 1J–L; Table S1**).

Notably, daytime light intensity exerted a strong effect on dark-phase sleep-wake distribution in female mice. Dim daytime light was associated with increased nocturnal wakefulness, whereas brighter daytime light progressively increased NREM and REM sleep during the dark phase (**Fig.1J–L**). In contrast, male mice exhibited minimal sensitivity to daytime light intensity. Together, these findings indicate that sex differences in sleep-wake architecture are driven primarily by increased nocturnal wakefulness in females under dim daytime light conditions.

To determine whether daytime light intensity affected sleep quality, we analyzed episode counts and durations for each vigilance state (**Fig. 2; Table S2**). A main effect of sex was observed for wake episode counts, with males exhibiting more wake episodes, consistent with more fragmented wakefulness compared with females (**Fig. 2A**). For NREM and REM sleep, episode counts were influenced primarily by the dark phase rather than daytime light intensity, with males exhibiting higher counts (**Fig. 2B**). A sex difference in the number of REM episodes was detected only under 50 lux daytime light condition (**Fig. 2C**). No significant effects of daytime light intensity were detected on episode counts for any vigilance state (**Fig. 2A–C**). Similarly, wake episode duration differed by sex, with females exhibiting longer wake bouts than males (**Fig. 2D**), whereas NREM and REM episode durations were not significantly affected by sex or daytime light intensity (**Fig. 2E, F**). Together these data indicate that female mice exhibit more consolidated wakefulness and sleep than male mice and vigilance state episode consolidation is largely independent of daytime light intensity.

**Figure 2.**
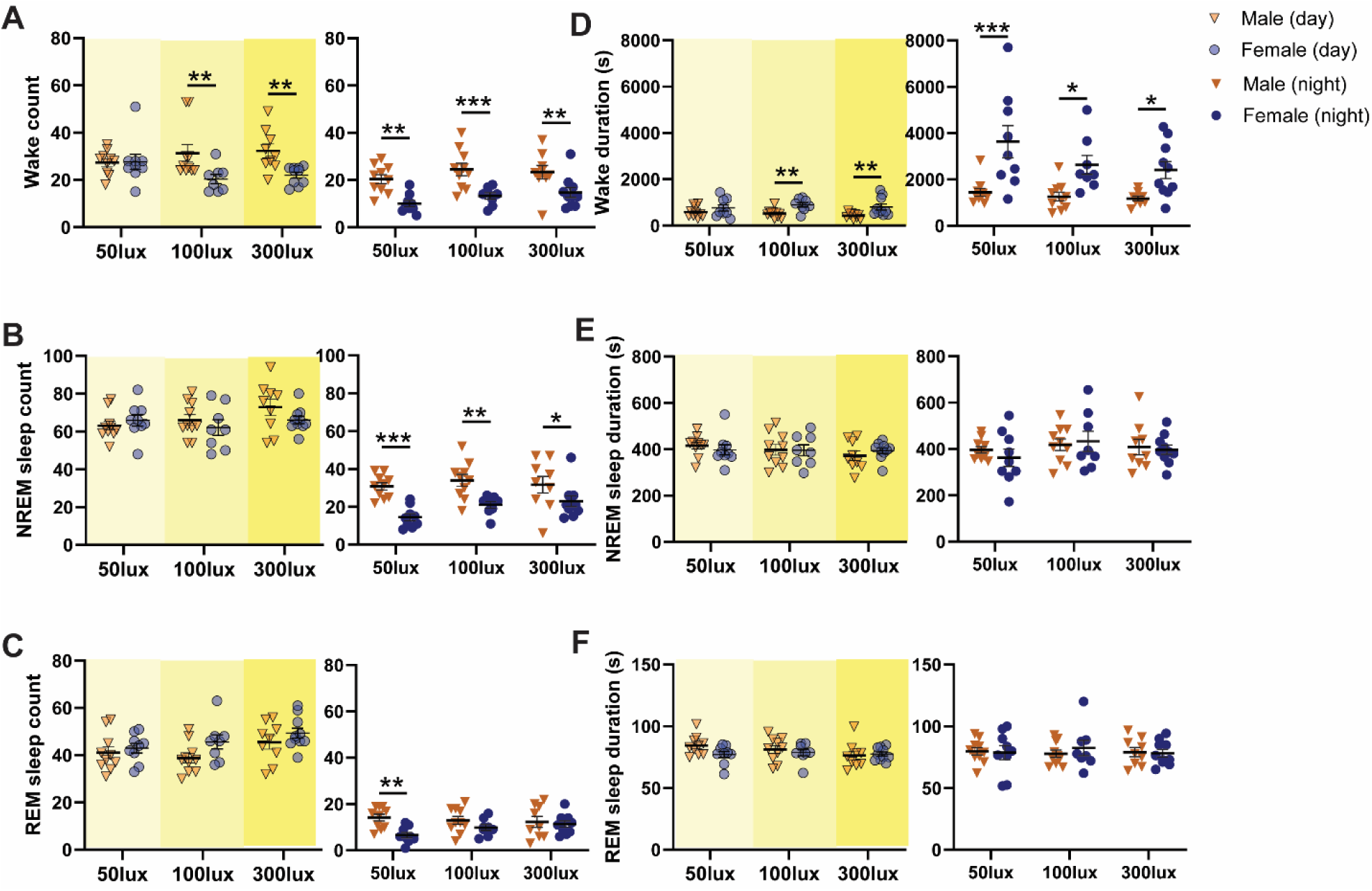
Bout number, duration, and state transitions in male and female WT mice under different daytime light intensities. (**A–C**) Number of bouts of wakefulness, NREM sleep, and REM sleep during the day (left) and night (right) in male and female mice exposed to 50, 100, or 300 lux daytime light. (**D–F**) Episode duration of wakefulness, NREM sleep, and REM sleep during the day (left) and night (right) in male and female mice exposed to 50, 100, or 300 lux daytime light. Asterisks indicate significant differences across daytime light intensity or between sexes (Bonferroni multiple-comparisons test after two-way ANOVA; * *p* < 0.05, ** *p* < 0.01, *** *p* < 0.001). Different shades of yellow represent different daytime light intensities. Data are presented as mean ± SEM. Sample sizes: 50 lux, 10 males and 9 females; 100 lux, 10 males and 8 females; 300 lux, 9 males and 10 females.

### Effects of dim light at night on sleep–wake architecture under different daytime light intensities

To determine whether daytime light intensity modulates the effects of dim light at night (DLaN) on sleep-wake architecture, we assessed sleep patterns under multiple daytime lighting conditions. In prior work, exposure to 5 lux DLaN for two weeks did not significantly alter sleep-wake architecture, suggesting that vulnerability to DLaN may depend on lighting context. We therefore increased nighttime illumination to 10 lux and examined both acute (one night) and chronic (two weeks) DLaN exposure. This design enabled direct comparison of sleep–wake architecture following acute and sustained DLaN across a range of daytime light intensities. Sleep architecture was evaluated over a 24-hour period in male and female mice housed under dim (50 lux), moderate (100 lux), or bright (300 lux) daytime light levels.

### Acute effects of dim light at night on sleep-wake architecture under different daytime light intensities

As anticipated, the effects of the first night of DLaN were strongly dependent on the daytime light intensity (**Fig. 3**). Mice housed under 50 lux 100 lux daytime lighting exhibited pronounced changes in sleep-wake architecture, whereas those housed under 300 lux daytime light showed relatively modest alterations. This pattern was observed in both sexes, although the magnitude and structure of the response differed between males and females.

**Figure 3.**
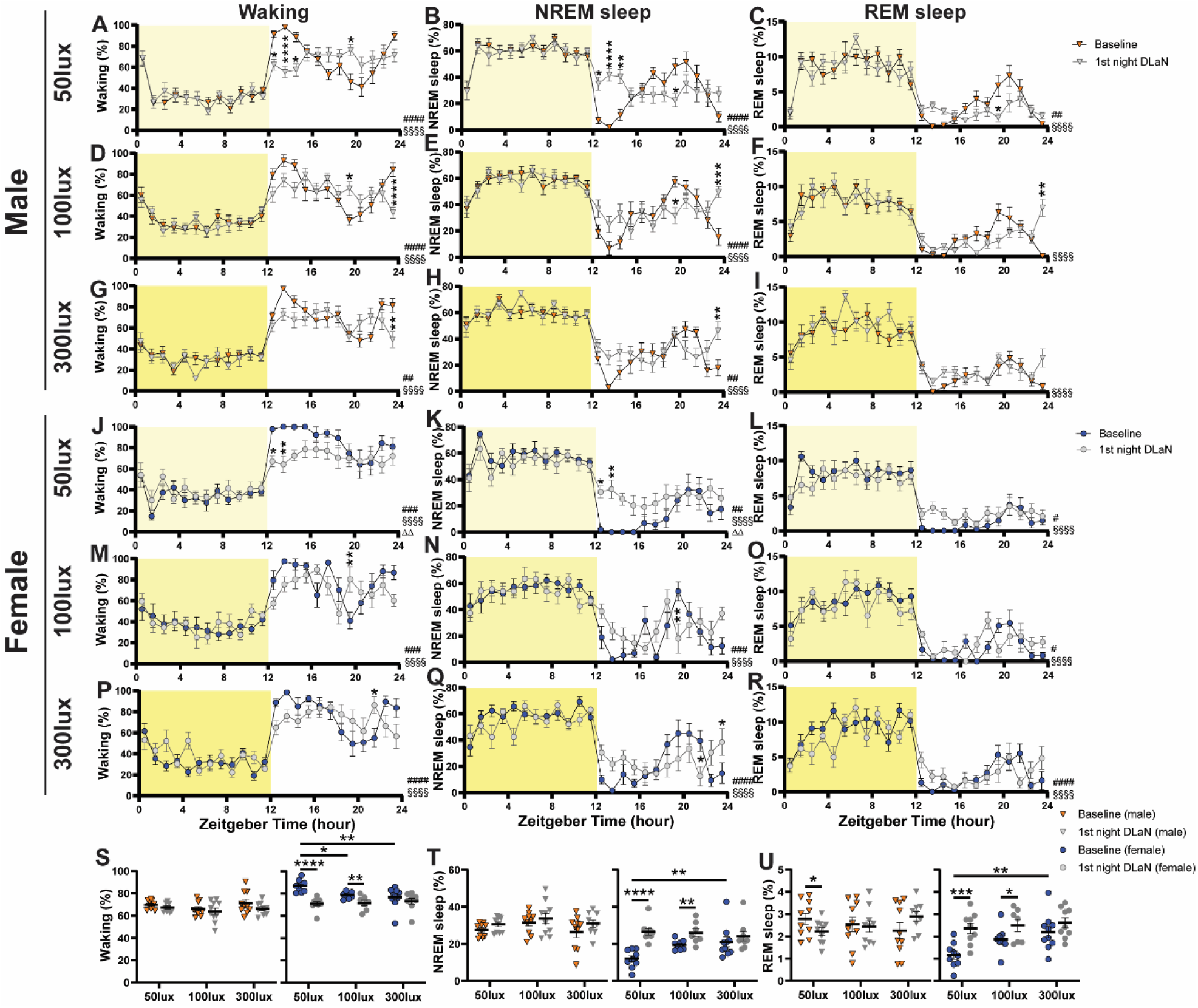
Sleep architecture of male and female WT mice during the first night of DLaN under different daytime light intensities. (**A–I**) 24-hour distribution of time spent in wakefulness (**A, D, G**), NREM sleep (**B, E, H**), and REM sleep (**C, F, I**) in male mice under baseline conditions (orange triangles) and during the first night of DLaN (gray triangles) following exposure to 50, 100, or 300 lux daytime light. Pound signs (#) indicate a significant interaction between zeitgeber time and first-night DLaN (two-way repeated-measures ANOVA or mixed-effects model with Geisser–Greenhouse correction; ## *p* < 0.01, #### *p* < 0.0001). Main effects of zeitgeber time are indicated by § (§§§§ *p* < 0.0001). Asterisks denote significant post hoc differences between baseline and first-night DLaN (Bonferroni multiple-comparisons test; * *p* < 0.05, ** *p* < 0.01, *** *p* < 0.001, **** *p* < 0.0001). (**J–R**) 24-hour distribution of time spent in wakefulness (**J, M, P**), NREM sleep (**K, N, Q),** and REM sleep (**L, O, R**) in female mice under baseline conditions (blue circles) and during the first night of DLaN (gray circles) following exposure to 50, 100, or 300 lux daytime light. Pound signs (#) indicate a significant interaction between zeitgeber time and first night DLaN (two-way repeated-measures ANOVA or mixed-effects model with Geisser–Greenhouse correction; # *p* < 0.05, ## *p* < 0.01, ### *p* < 0.001, #### *p* < 0.0001). Main effects of zeitgeber time and first-night DLaN are indicated by § and Δ, respectively (§§§§ *p* < 0.0001; ΔΔ *p* < 0.01). Asterisks denote significant post hoc differences between baseline and first night DLaN (Bonferroni multiple-comparisons test; * *p* < 0.05, ** *p* < 0.01). (**S–U**) Nighttime distribution of time spent in wakefulness, NREM sleep, and REM sleep under baseline conditions and during the first night of DLaN in male (left, triangle) and female mice (right, circle) under different daytime light intensities. Asterisks indicate significant post hoc comparisons (Bonferroni multiple-comparisons test; * *p* < 0.05, ** *p* < 0.01, *** *p* < 0.001). Different shades of yellow represent different daytime light intensities. Sample sizes: 50 lux, 10 males and 9 females; 100 lux, 10 males and 8 females (first-night DLaN: 7 females); 300 lux, 10 males and 10 females (first-night DLaN: 9 males and 9 females). Data are presented as mean ± SEM.

In males, acute DLaN exposure induced time-specific changes that varied with daytime light intensity. Under 50 lux daytime light, males exhibited reduced wakefulness and increased NREM sleep during the early dim light phase (ZT13–15), followed by increased wakefulness and reduced REM sleep later in the night (ZT20; **Fig. 3A–C**). Under 100 lux, males showed increased wakefulness and reduced NREM sleep at ZT20, followed by increased sleep at the end of the dim light phase (ZT24; **Fig. 3D–F**). Under 300 lux daytime light, effects were limited to a modest reduction in wakefulness and increase in NREM sleep at ZT24 (**Fig. 3G–I**). Females displayed a distinct and more pronounced response to acute DLaN exposure. Under 50 lux daytime light, females showed reduced wakefulness and increased NREM sleep early in the dim light phase (ZT13–14; **Fig. 3J–L**). Under 100 lux, females exhibited increased wakefulness and reduced NREM sleep at ZT20 (**Fig. 3M–O**). Under 300 lux, changes were modest and restricted to late-night time points, with increased wakefulness at ZT22 and increased NREM sleep at ZT24 (**Fig. 3P–R**).

When sleep-wake states were quantified across the entire 12-hour dim light phase, first-night DLaN had a main effect on nighttime wakefulness and NREM sleep in males, but no main effect of daytime light intensity was detected (**Fig. 3S, T**). However, a significant interaction between first night DLaN exposure and daytime light intensity was observed for REM sleep (**Fig. 3U**). Specifically, under 50 lux daytime light, male mice exhibited a modest but significant reduction in REM sleep following the first night of DLaN exposure. In contrast, females exhibited robust sensitivity to both daytime light intensity and acute DLaN exposure. Significant interactions between DLaN exposure and daytime light intensity were observed for wakefulness and NREM sleep, along with a main effect of DLaN across all vigilance states (**Fig. 3S–U; Table S3**). Under both 50 and 100 lux daytime light, acute DLaN exposure markedly reduced wakefulness and increased both NREM and REM sleep across the dim light phase, effects that were not observed under 300 lux daytime light.

Because light at night is commonly associated with sleep fragmentation, we next examined sleep-wake episode structure following acute DLaN exposure. A single night of DLaN increased bout counts for both wakefulness and sleep states in males and females, indicating disrupted sleep–wake continuity independent of sex and daytime light intensity (**Supplemental Fig. 1**). Together, these findings demonstrate that daytime light intensity gates acute sensitivity to light at night, with females exhibiting heightened vulnerability under dim and moderate daytime lighting conditions.

### Chronic effects of dim light at night on sleep–wake architecture under different daytime light intensities

Because the effects of acute DLaN were highly dependent on daytime light intensity and differed by sex, we next examined whether these responses persist, adapt, or worsen following chronic DLaN exposure. After 2 weeks of DLaN exposure, sleep-wake alterations became more pronounced and diverged markedly from the acute responses observed after a single night of DLaN. As with acute exposure, chronic DLaN did not significantly affect sleep-wake organization during the light phase; instead, the most prominent alterations occurred during the dim light phase (**Fig. 4**).

**Figure 4.**
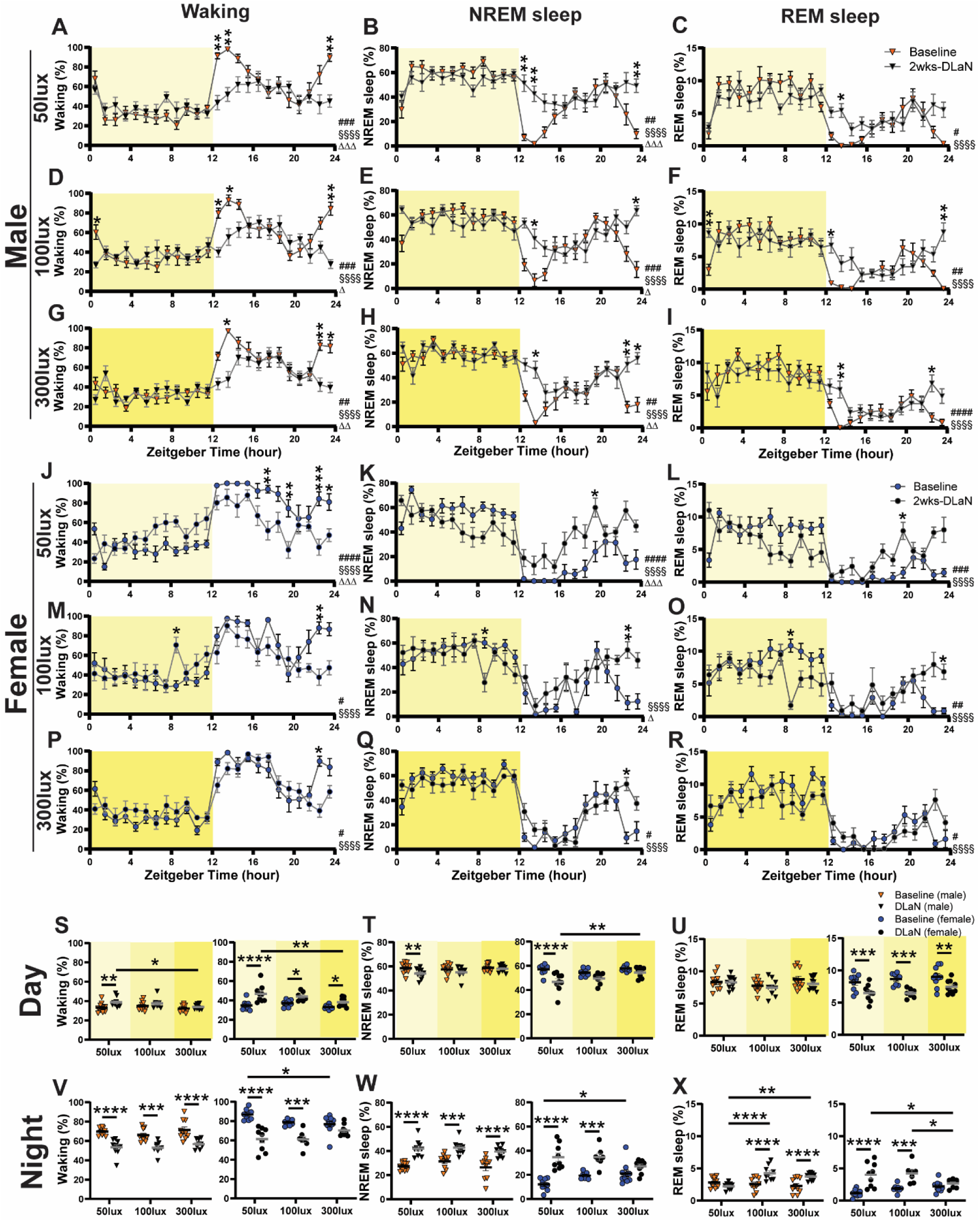
Sleep architecture of male and female WT mice following 2 weeks of DLaN under different daytime light intensities. (**A–I**) 24-hour distribution of time spent in wakefulness (**A, D, G**), NREM sleep (**B, E, H**), and REM sleep (**C, F, I**) in male mice under baseline conditions (orange triangles) and after 2 weeks of DLaN exposure (black triangles) following exposure to 50, 100, or 300 lux daytime light. Pound signs (#) indicate a significant interaction between zeitgeber time and DLaN (two-way repeated-measures ANOVA or mixed-effects model with Geisser–Greenhouse correction; # *p* < 0.05, ## *p* < 0.01, ### *p* < 0.001, #### *p* < 0.0001). Main effects of zeitgeber time and DLaN are indicated by § and Δ, respectively (§§§§ *p* < 0.0001; Δ *p* < 0.05, ΔΔ *p* < 0.01, ΔΔΔ *p* < 0.001). Asterisks denote significant post hoc differences between baseline and DLaN (Bonferroni multiple-comparisons test; * *p* < 0.05, ** *p* < 0.01). (**J–R**) 24-hour distribution of time spent in wakefulness (**J, M, P**), NREM sleep (**K, N, Q),** and REM sleep (**L, O, R**) in female mice under baseline conditions (blue circles) and after 2 weeks of DLaN exposure (black circles) following exposure to 50, 100, or 300 lux daytime light. Pound signs (#) indicate a significant interaction between zeitgeber time and DLaN (two-way repeated measures ANOVA or mixed-effects model with Geisser–Greenhouse correction; # *p* < 0.05, ## *p* < 0.01, ### *p* < 0.001, #### *p* < 0.0001). Main effects of zeitgeber time and DLaN are indicated by § and Δ, respectively (§§§§ *p* < 0.0001; Δ *p* < 0.05, ΔΔΔ *p* < 0.001). Asterisks denote significant post hoc differences between baseline and DLaN (Bonferroni multiple-comparisons test; * *p* < 0.05, ** *p* < 0.01, *** *p* < 0.001). (**S–X**) Daytime and nighttime distribution of time spent in wakefulness, NREM sleep, and REM sleep under baseline conditions and after 2 weeks of DLaN exposure in male (left, triangle) and female mice (right, circle) under different daytime light intensities. Asterisks indicate significant post hoc comparisons (Bonferroni multiple-comparisons test; * *p* < 0.05, ** *p* < 0.01, *** *p* < 0.001, **** *p* < 0.0001). Different shades of yellow represent different daytime light intensities. Sample sizes: 50 lux, 10 males and 9 females; 100 lux, 10 males and 8 females (DLaN: 7 females); 300 lux, 10 males and 10 females (DLaN: 9 males and 9 females). Data are presented as mean ± SEM.

In male mice, chronic DLaN elicited a highly consistent and reproducible response across all three daytime light intensities. Compared with baseline LD conditions, males exhibited increased sleep and reduced wakefulness at both the beginning and end of the dim light phase under 50, 100, and 300 lux daytime light (**Fig. 4A–I**). Significant interactions between zeitgeber time and DLaN exposure were detected for wakefulness, NREM sleep, and REM sleep across all daytime light conditions (**Table S4**), indicating a robust and light-intensity-independent response to chronic DLaN. This response pattern closely mirrors previous reports of chronic DLaN exposure in male rodents, including a 14-day DLaN paradigm in rats.^16^

In contrast, female mice exhibited a strikingly different and unexpected response to chronic DLaN exposure. Unlike males, females did not show reduced wakefulness or increased NREM sleep at the onset of the dim light phase, with no significant differences observed at ZT13 or ZT14 under any daytime light condition (**Fig. 4J–R**). Instead, the temporal structure of the DLaN response in females depended strongly on daytime light intensity. Under 50 lux daytime light, wakefulness was reduced during the middle and end of the dim light phase (ZT18, 20, 23, 24), accompanied by increases in both NREM and REM sleep. Under 100 and 300 lux daytime light condition, reductions in wakefulness and increases in NREM sleep were largely restricted to the end of the dim light phase (ZT23; **Fig. 4J–R**). Notably, unlike male mice, females also exhibited DLaN-induced alterations during the light phase. Under 100 lux daytime light, chronic DLaN increased wakefulness and reduced sleep during the mid-light phase (ZT 9; **Fig.4M–O**). Significant interactions between time and DLaN exposure were detected across all three daytime light intensities in females (**Table S4**), indicating a temporally dynamic but context-dependent response to chronic DLaN. Regardless of sex, both sexes exhibited a similar anticipatory pattern prior to lights on (ZT 0) under baseline LD conditions, characterized by increased wakefulness, consistent with anticipation of the light phase. Across all light conditions, this increase in wakefulness was markedly attenuated under chronic DLaN exposure, indicating a loss of anticipatory of the start of light phase in both sexes.

To assess total amount of sleep and wakefulness, we next quantified total 24-hour amounts of vigilance states under baseline and chronic DLaN conditions. Male mice displayed a consistent response across all three daytime light intensities, characterized by decreased wakefulness and increased NREM sleep, with relatively minor effects on REM sleep (**Supplemental Fig.2 A-C**). The female mice housed under 50 and 100 lux daytime light conditions also showed similar overall pattern, whereas females housed under 300 lux daytime light conditions did not exhibit significant changes in total wakefulness or NREM sleep. In contrast to males, REM sleep responses in females was strongly modulated by daytime light intensity, increasing under 50 lux but decreasing under 300 lux conditions (**Supplemental Fig.2 D-F**).

Because the largest effects were observed during the dark phase, we further analyzed sleep–wake distribution separately for the light and dark/dim light phases. During the light phase, male mice exhibited modest increases in wakefulness and corresponding decreases in NREM sleep under 50 lux daytime light, with no significant effects on REM sleep under any lighting condition (**Fig. 4S–U**). These limited light-phase effects in males are consistent with prior DLaN studies. Female mice, however, showed robust light-phase responses to chronic DLaN across all daytime light intensities, characterized by increased wakefulness and reduced REM sleep (**Fig. 4S, U**). In contrast, dark-phase responses were more similar between sexes. Male mice showed decreased wakefulness and increased NREM sleep across all three daytime light conditions, whereas females exhibited comparable dim light-phase changes under 50 and 100 lux but not under 300 lux daytime light (**Fig. 4V–X**). The absence of significant dim light-phase effects in females under bright daytime light suggests that higher daytime illumination partially protects against DLaN-induced sleep disruption.

### Effect of dim light at night on quality of sleep and wakefulness under different daytime light intensities

We next examined sleep-wake fragmentation following two weeks of DLaN exposure. Consistent with the acute effects observed after the first night of DLaN, bout counts during the dim light phase were significantly increased, indicating that DLaN induces persistent fragmentation of sleep and wakefulness (**Supplemental Fig.3**). The similarity between acute and chronic exposure suggests that disruption of sleep continuity emerges rapidly and is maintained with prolonged DLaN exposure. In contrast, the light phase effects of DLaN differed markedly by sex. In male mice, bout counts during the light phase were not significantly altered. However, in female mice, chronic DLaN exposure led to reduced bout counts during the light phase, particularly for REM and NREM sleep under 50 lux daytime light and increased wake bout counts under 300 lux daytime light, indicating a sex-specific sensitively to DLaN during the light phase.

Analysis of episode duration revealed that chronic DLaN selectively shortened wake episode duration during the dim light phase, whereas NREM and REM sleep episode durations were not significantly affected. Finally, analysis of vigilance state transitions found sex-specific and phase-dependent differences in transition patterns during both the light and dark/dim light phases, indicating that DLaN reshapes sleep–wake dynamics beyond simple changes in state occupancy (**Supplemental Fig.4**).

### Effects of acute and chronic dim light at night on electroencephalogram slow wave activity in NREM sleep in male and female mice under different daytime light intensities

During the first night of DLaN exposure (**Fig. 5A-F**), neither male nor female mice exhibited a significant interaction between DLaN and ZT, indicating minimal acute effects on NREM sleep electroencephalogram (EEG) slow wave activity (SWA, EEG power density between 0.5-4.0 Hz, an important marker for quality and depth of sleep; **Table S5**). In male mice housed under 100 lux daytime light, acute DLaN exposure was associated with transient increases in SWA at late dark-phase time points (ZT 21, 23). In contrast, under 300 lux daytime light, no significant main effects or interactions with DLaN were observed in either sex. Together these findings suggest that brighter daytime light attenuates acute DLaN-induced alterations in SWA across sexes.

**Figure 5.**
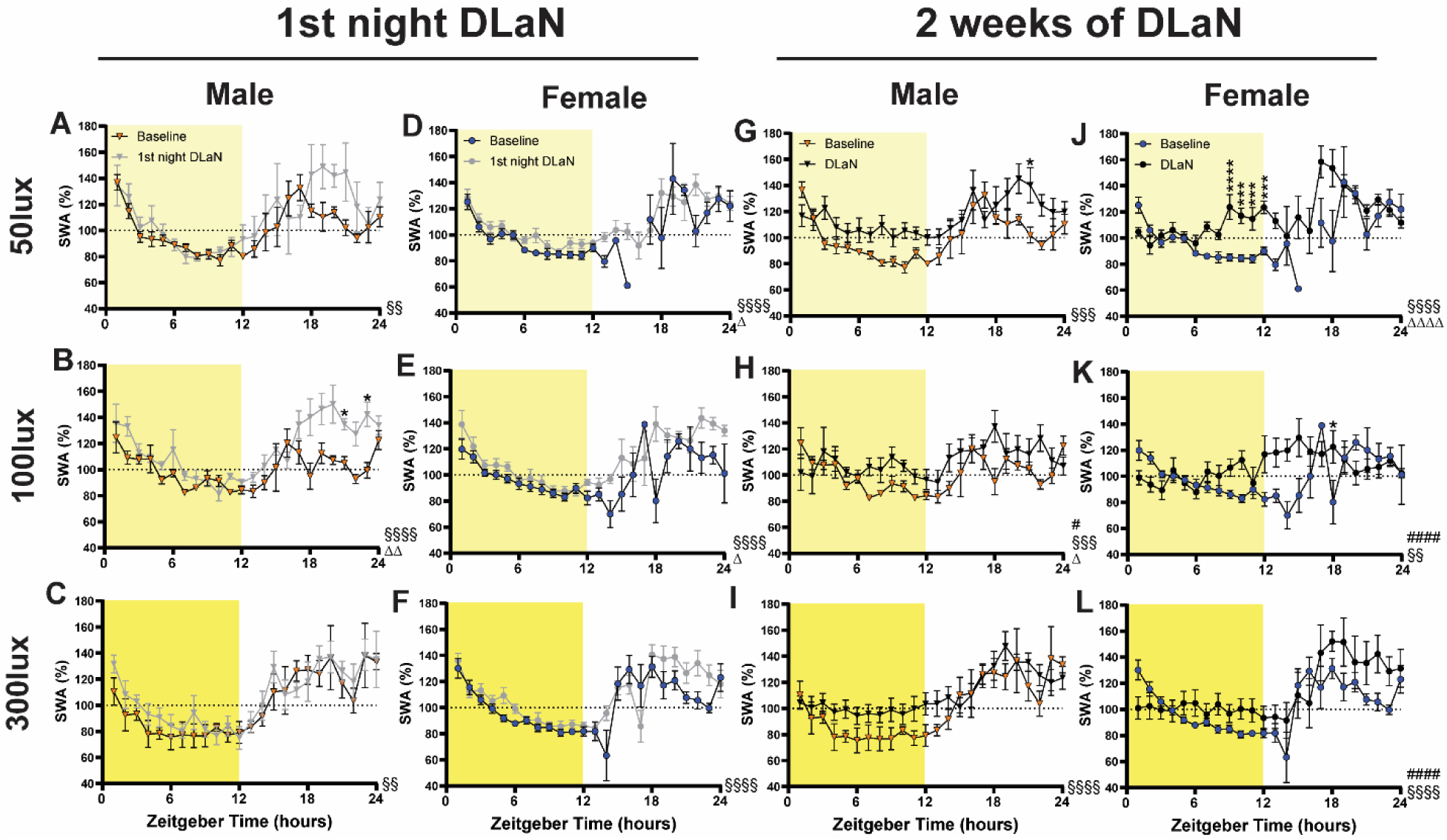
NREM sleep slow-wave activity in male and female WT mice exposed to 1^st^ night of DLaN and 2 weeks of DLaN under different daytime light intensities. (**A–F**) 24-hour distribution of NREM sleep slow-wave activity in male (**A-C**; triangles) and female (**D-F**; circles) mice following exposure to 50, 100, or 300 lux daytime light, during baseline (orange or blue) and the first night of DLaN (gray). (**G–L**) 24-hour distribution of NREM sleep slow-wave activity in male (**G-I**; triangles) and female (**J-L**; circles) mice following exposure to 50, 100, or 300 lux daytime light, during baseline and after 2 weeks of DLaN exposure (black). Due to missing time points in the 50lux female baseline, only main effects were analyzed in panels D and J; post hoc analyses were performed using a mixed-effects model excluding data from ZT13–ZT16. Pound signs (#) indicate a significant interaction between zeitgeber time and DLaN (mixed-effects model with Geisser–Greenhouse correction; # *p* < 0.05, ## *p* < 0.01, #### *p* < 0.0001). Main effects of zeitgeber time and DLaN are indicated by § and Δ, respectively ((§§ *p* < 0.01, §§§ *p* < 0.001, §§§§ *p* < 0.0001; Δ *p* < 0.05, ΔΔ *p* < 0.01, ΔΔΔΔ *p* < 0.0001). Asterisks denote significant post hoc differences between baseline and DLaN conditions (Bonferroni multiple-comparisons test; * *p* < 0.05, *** *p* < 0.001, **** *p* < 0.0001). Different shades of yellow represent different daytime light intensities. Data are presented as mean ± SEM.

Following two weeks of DLaN exposure, sex- and light intensity-dependent differences in NREM SWA emerged. In male mice, a significant interaction between DLaN exposure and ZT was observed under 100 lux daytime light (**Fig. 5H**), whereas no significant interaction or main effect of DLaN was detected under 50 or 300 lux conditions (**Figure 5G, I**). Notably, under 50 lux daytime light, males exhibited a localized increase in SWA at ZT20 (**Fig. 5G**). In contrast, female mice exhibited more robust and widespread SWA responses to chronic DLaN exposure across all daytime light intensities. Under 50 lux daytime light, females showed a pronounced increase in SWA during the light phase, particularly between ZT 9 and ZT 12 (**Fig. 5J**). Significant interactions between DLaN exposure and ZT were also observed under both 100 lux and 300 lux daytime light conditions (**Fig. 5K**, **L**), indicating temporally reorganized SWA dynamics rather than isolated changes in magnitude.

Power spectral analysis of NREM sleep further supported these sex-specific effects. In male mice, chronic DLaN exposure produced a significant increase in delta power. In female mice, a significant main effect of DLaN on NREM spectral power was observed, although no frequency-specific changes were detected (**Supplemental Fig. 5**), consistent with a redistribution of SWA timing rather than a selective enhancement of delta-band activity.

### Circadian timing and amplitude of the sleep–wake rhythm under baseline and DLaN conditions

As slow wave activity during NREM sleep was strongly influenced by the endogenous circadian clock, and in our female mice, we observed a strong interaction between time and DLaN. We next examined whether the daily timing and amplitude of vigilance states were altered under DLaN conditions. To address this, we applied JTK_CYCLE analysis to baseline and DLaN datasets, allowing us to assess changes in the circadian peak time and amplitude of wakefulness, NREM sleep, and REM sleep under different daytime light intensities.

Surprisingly, male mice did not exhibit any significant changes in the peak timing of wakefulness, NREM sleep, or REM sleep under DLaN conditions across all daytime light intensities (**Fig. 6A–I**). In contrast, female mice showed significant phase advances compared with baseline under dimmer daytime light conditions. Under 50 lux daytime light, females exhibited a marked phase advance around 4 hours in wakefulness, NREM sleep and REM sleep (**Fig. 6S, V, Y**). Similarly, under 100 lux daytime light conditions, female mice showed significant phase advances around 3 hours compared with baseline in wakefulness and NREM sleep (**Fig. 6T, W**). Thus, under both 50 and 100 lux conditions, a phase advance of approximately 3–4 hours emerged as a consistent feature of the female response to chronic DLaN. This advance likely explains why female mice did not exhibit the expected decrease in wakefulness or increase in sleep at the beginning of the active phase after two weeks of DLaN exposure: their peak in waking had shifted to earlier ZT. Notably, under 300 lux daytime light conditions, neither the timing of the vigilance state peaks nor overall sleep–wake amounts were significantly altered, suggesting that brighter daytime light may protect against DLaN induced phase shifts and their downstream effects on sleep architecture in female mice.

**Figure 6.**
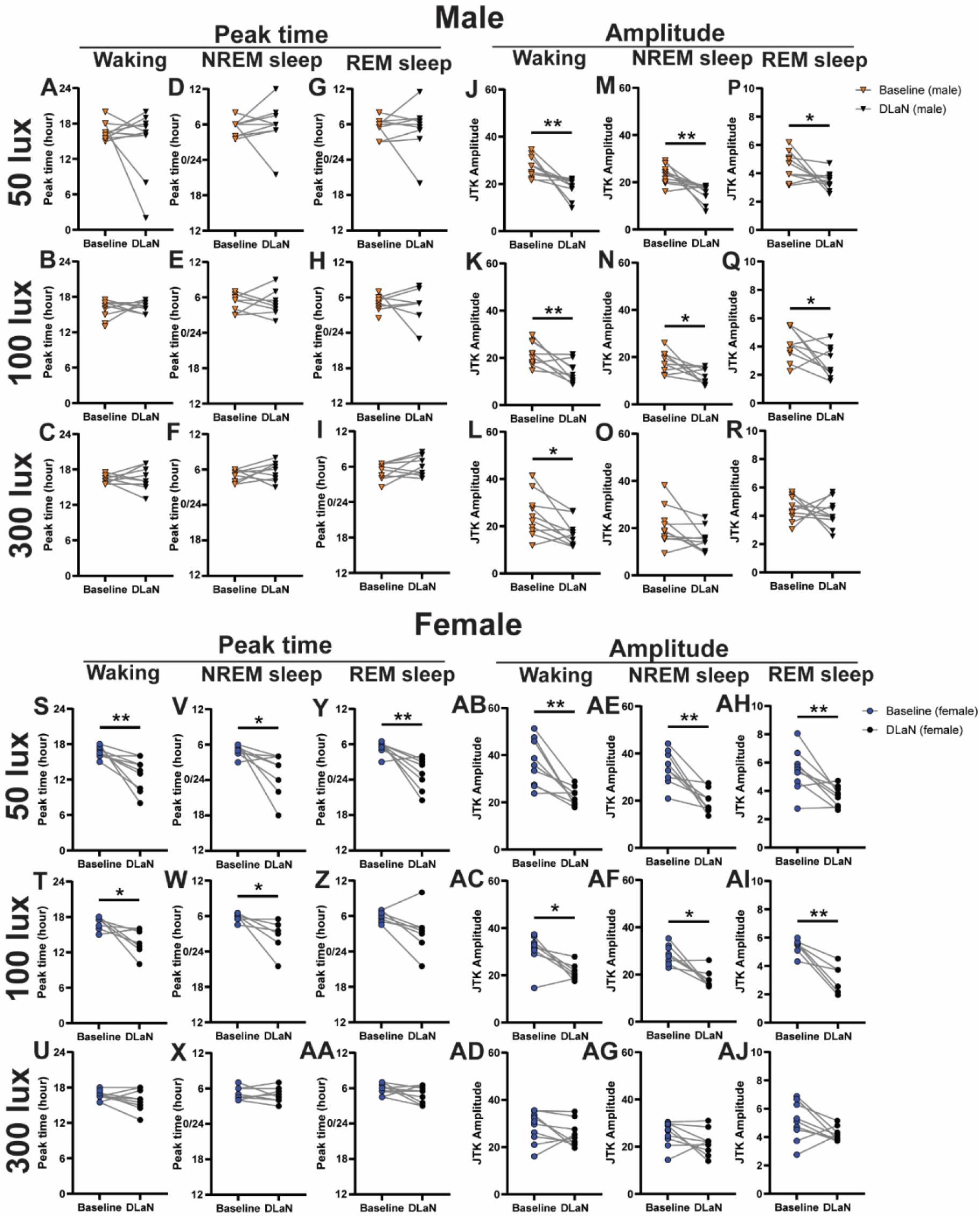
Peak timing and amplitude of wakefulness, NREM and REM sleep in male and female WT mice following 2 weeks of DLaN under different daytime light intensities. **(A–I)** Peak timing of wakefulness (**A–C**), NREM sleep (**D–F**), and REM sleep (**G–I**), identified using JTK-Cycle, in male mice exposed to 50 lux (**A, D, G**), 100 lux (**B, E, H**), or 300 lux (**C, F, I**) daytime light under baseline conditions (orange triangles) and after 2 weeks of DLaN exposure (black triangles). **(J–R)** Amplitude of wakefulness (**J–L**), NREM sleep (**M–O**), and REM sleep (**P–R**), identified using JTK-Cycle, in male mice exposed to 50 lux (**J, M, P**), 100 lux (**K, N, Q**), or 300 lux (**L, O, R**) daytime light under baseline conditions (orange triangles) and after 2 weeks of DLaN exposure (black triangles). **(S–AA)** Peak timing of wakefulness (**S–U**), NREM sleep (**V–X**), and REM sleep (**Y–AA**), identified using JTK-Cycle, in female mice exposed to 50 lux (**S, V, Y**), 100 lux (**T, W, Z**), or 300 lux (**U, X, AA**) daytime light under baseline conditions (blue circles) and after 2 weeks of DLaN exposure (black circles). **(AB–AJ)** Amplitude of wakefulness (**AB–AD**), NREM sleep (**AE–AG**), and REM sleep (**AH–AJ**), identified using JTK-Cycle, in female mice exposed to 50 lux (**AB, AE, AH**), 100 lux (**AC, AF, AI),** or 300 lux (**AD, AG, AJ**) daytime light under baseline conditions (blue circles) and after 2 weeks of DLaN exposure (black circles). Asterisks indicate significant differences assessed by paired *t*-tests or Wilcoxon matched pairs signed-rank tests (* *p* < 0.05, ** *p* < 0.01). Sample sizes: 50 lux, 10 males and 9 females; 100 lux, 10 males and 8 females (DLaN: 7 females); 300 lux, 10 males and 10 females (DLaN: 9 males and 9 females). Data are shown as individual values.

We next analyzed the amplitude of daily oscillations in vigilance states. In male mice, DLaN exposure led to significant reductions in oscillation amplitude for wakefulness (**Fig. 6J–L**), NREM sleep (**Fig. 6M–N**), and REM sleep (**Fig. 6P–Q**) across all conditions. In female mice, decreases in amplitude were observed for wakefulness (**Fig. 6AB–AC**), NREM sleep (**Fig. 6AE–AF**), and REM sleep (**Fig. 6AH–AI**) under 50 and 100 lux daytime light conditions. However, female mice housed under 300 lux daytime light did not exhibit significant amplitude reductions, further supporting the notion that brighter daytime light may protect against DLaN induced disruptions in both the timing and strength of daily sleep–wake rhythms, particularly in females. Together these data show that chronic DLaN differentially affects timing and amplitude of sleep-wake rhythms in males and females with females showing more susceptibility to DLaN.

## Methods

### Animals and housing conditions

11 weeks old C57BL/6J wild-type mice (30 female, 30 male) and were used in this study. All animal procedures were performed in accordance with the UCLA animal care committee’s regulations. C57BL/6J wild-type mice were obtained from the Jackson Laboratory (ME, USA). Mice were housed in light-tight ventilated cabinets in temperature- and humidity-controlled conditions, with free access to food and water.

### Surgical procedure

At the age of 12 weeks, animals were anesthetized using ketamine/xylazine (100/8.8 mg/kg, intraperitoneal injection) and EEG/ EMG electrodes were implanted as described previously.^4^ A prefabricated head mount (CAT:8201, Pinnacle Technologies, KS) was used to position four stainless-steel epidural screw electrodes. One front electrode was placed 1.5 mm anterior to bregma and 1.5 mm lateral to the central line, whereas the second two electrodes (interparietal—located over the visual cortex and common reference) were placed 2.5 mm posterior to bregma and 1.5 mm on either side. A fourth screw served as a ground. Electrical continuity between the screw electrode and head mount was aided by silver epoxy. EMG activity was monitored using stainless-steel Teflon-coated wires that were inserted bilaterally into the nuchal muscle. The head mount (integrated 2 × 3 pin grid array) was secured to the skull with dental cement.

### Experimental design and recording setup

60 mice were randomly divided into three groups, light intensity during the day was 50 lux, 100lux and 300lux as measured at the base of the cage, and 0 lx during the night (**Fig. 7A**). All mice were first entrained to a normal lighting cycle: 12 h light: 12 h dark (LD) before the surgery for 1 week. After the surgery, these mice recovered under these 3 different daylight conditions for two weeks, 3 female mice died during the surgery, two female mice from 100lux group and 1 female mouse from 50lux group. Mice were connected to a lightweight tether attached to a low-resistance commutator mounted over the cage (Pinnacle Technologies, KS) as previously described. Mice were allowed a minimum of 2 additional days to acclimate to the tether and recording chamber. Subsequently, a baseline day was recorded, starting at lights on zeitgeber time 0. After the baseline recording, mice were exposure to a 10lux dim light at night for one day, was recorded as 1^st^ night DLaN. And these mice were continued with two weeks of DLaN, the mice were attached to the commutator for the post 2 weeks DLaN recording for 24 hrs (**Fig. 7B**).

**Figure 7.**
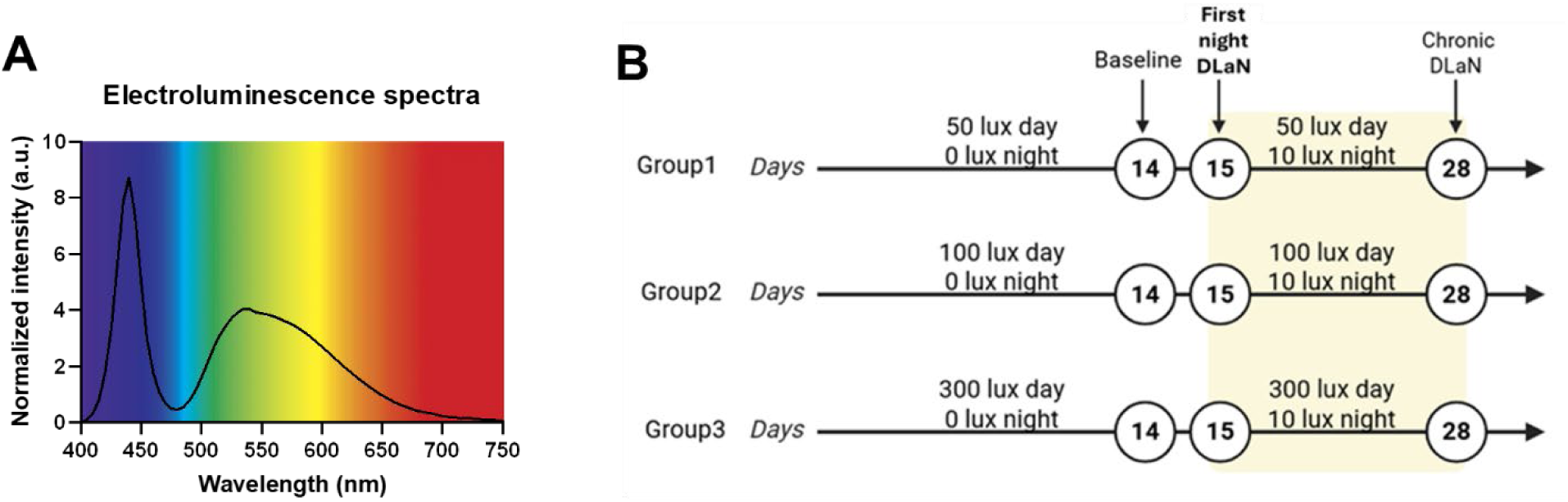
LED spectra and experimental design. A. The normalized electroluminescence spectra (a.u., arbitrary unit) of white LED sources. B. Detailed experimental design

### EEG data acquisition, sleep scoring and EEG power spectrum analysis

Data acquisition was done by Sirenia Acquisition software (Pinnacle Technologies, KS) as previously described. EEG signals were low pass filtered with a 40 Hz cutoff and collected continuously at a sampling rate of 400 Hz. All data were recorded simultaneously in 1s epoch. Wake, NREM, and REM sleep were determined visually. Wake was scored when there were a high-EMG and low-EEG amplitude and high theta activity (EEG power density in the theta band, 6.0–9.0 Hz), concomitant with irregular, high EMG values. NREM sleep was scored when there was a low-EMG and higher EEG amplitude compared with wake and high SWA. REM sleep was scored when the EMG and EEG amplitude was low, and high theta activity was visible in the EEG.

Movement artifacts were excluded for power spectral and slow-wave analysis. For the spectral analysis, complete and clean recordings are needed to enter the analysis. However, this was not possible in six animals, which were therefore excluded from the analysis of the absolute EEG power density spectra and SWA in NREM sleep. Spectral analysis was performed using fast Fourier transform (FFT; 0.1–40 Hz, 0.1 Hz resolution); the absolute power density spectra of NREM sleep in baseline and post DLaN days were analyzed.

Three vigilance states (wake, NREM, and REM sleep) were scored offline in 10-s epochs by the Sirenia program. The manual scoring of vigilance states based on the EEG and EMG recordings was performed according to standardized criteria for mice. The average amount of the vigilance states (wake, NREM sleep, and REM sleep) and EEG spectral data and NREM sleep SWA were analyzed in 1 hr, 12 hr and 24 hr intervals.

### Bouts, episode duration and transitions analysis

Episode duration averages were compiled by the Sirenia program, which uses a conservative algorithm for bout lengths requiring sustained changes in arousal state to switch state. A bout was defined as 3+ continuous epochs, and the occurrence of 3+ continuous epochs of the different stage indicated the end of the current bout. Episode duration and bout number were analyzed in 12 hr intervals. To evaluate state transitions, we used custom-written Pascal scripts to determine transitions between different vigilance states. A transition was included in the analysis when immediately before the transition at least 30 consecutive seconds (3 scored epochs) were of the corresponding first vigilances state, and immediately after the transition also at least 30 consecutive seconds of the second correspond vigilance state were found.

### Spectral EEG analysis

To investigate the effect of DLaN on EEG power density of NREM sleep, we analyzed the relative EEG power density in the slow-wave range (SWA, 0.5–4.0 Hz) in NREM sleep as described previously.^4^ Since for the slow wave activity analysis, it was necessary to re-calculate power density values relative to the control condition, complete and clean recordings were needed from all animals for both conditions to enter the analysis. Unfortunately, this was not possible in 3 male animals under 2 weeks of DLaN, which were therefore excluded from the analysis of SWA.

### Peak timing and amplitude analysis

Circadian peak timing and amplitude of vigilance states were analyzed using JTK_CYCLE, an efficient nonparametric algorithm for detecting rhythmic components in time-series datasets. This approach was applied to determine peak time and amplitude parameters for each vigilance state, allowing within-animal comparisons between baseline and DLaN conditions. Analyses were performed using Nitecap.^17,18^

### Statistical analysis

All statistical analyses were conducted using GraphPad Prism 10 (GraphPad Software). Hourly (1-h) data in **Fig. 1** were analyzed using either mixed-effects models (REML) or two-way repeated-measures ANOVA (repeated zeitgeber time) with the Geisser–Greenhouse correction. Hourly (1-h) data in **Fig. 3-5** were analyzed using either mixed-effects models (REML) or two-way repeated-measures ANOVA (repeated in zeitgeber time and repeated measures under 1^st^ night DLaN or DLaN) with the Geisser–Greenhouse correction. Significant ANOVA results were followed by Bonferroni post hoc tests. For 12-h and 24-h summary data, two-way ANOVA or mixed-effects models (REML) were used to assess the effects of sex, DLaN exposure, or daytime light intensity. Normality was assessed using the Anderson–Darling test, or the Shapiro–Wilk test when sample size was less than eight. When data were normally distributed, paired Student’s t-tests were used; otherwise, the nonparametric Wilcoxon matched-pairs signed-rank test was applied to compare baseline and DLaN conditions for peak timing and amplitude. Detailed statistical results, including sample sizes for each panel, are provided in **Supplemental Tables 1–6.**

## Discussion

### Daytime light shapes sex differences in sleep-wake architecture

Light is a fundamental regulator of circadian and sleep–wake behavior, yet the influence of daytime light intensity has been studied primarily in humans and in male laboratory animals.^19^ In diurnal species, brighter daytime light strengthens circadian rhythms and improves sleep consolidation.^10^ In nocturnal rodents brighter daytime light has often been assumed to be detrimental, based largely on studies using constant illumination or extremely high light intensities (>1000 lux) in albino strains, conditions that are likely to induce retinal pathology rather than physiological circadian responses.^20,21^ In contrast, there is little evidence that moderately brighter daytime light impairs sleep or circadian regulation under stable light-dark conditions.

Whether commonly used daytime light intensities differentially affect sleep-wake regulation across sexes has remained unclear. Across the rodent literatures, reported sex differences in sleep-wake architecture are inconsistent, and daytime light intensity is rarely reported, complicating comparisons across species. We systematically examined sleep-wake architecture in male and female mice housed under 50, 100 and 300 lux daytime light intensity and found that sex differences in sleep-wake behavior depend critically on light intensity: female mice exhibited increased wakefulness under dim daytime light (50 and 100 lux), whereas no sex differences were observed under brighter daytime light (300 lux), consistent with our previous study conducted under the 300 lux conditions.^4^ These sex differences were driven primarily by increased wakefulness in females under dim daytime light rather than by changes in males, while sleep quality was largely preserved across all conditions, indicating that daytime light modulates the amount of sleep and wakefulness rather than its quality.

### Sex-specific resilience and vulnerability to light at night

Acute and chronic DLaN exert distinct effects on sleep-wake behavior that are strongly sex- and daytime light intensity-dependent. A single night of DLaN produced rapid but temporally restricted changes in sleep continuity and fragmentation in both sexes, indicating that even brief nighttime light exposure can disrupt sleep. These acute effects were broadly conserved: males and females showed transient increases in sleep, with subtle differences in magnitude and temporal distribution. In humans, similar single-night dim light exposure increases sleep duration, ^22,23^ though daytime light was not controlled. Prolonged DLaN, in contrast, induced more persistent alterations in sleep-wake architecture, including dampened daily amplitude, disrupted anticipation of lights-on, and sex-specific changes in the timing and distribution of vigilance states. Males maintained a largely conserved temporal structure with reduced circadian amplitude, exhibiting delayed wake onset and increased sleep at both the beginning and end of the dim light phase, similar patterns previously reported in rodents.^16,24^ Females, however, showed pronounced phase advances and redistribution of sleep and wakefulness across the 24-hour cycle, with reduced wakefulness and increased sleep late in the dim light phase but no delayed onset, revealing distinct strategies of circadian reorganization across sexes.

Both sexes displayed a loss of anticipatory wakefulness prior to light-on, a pattern reminiscent of human studies linking nighttime light exposure to delayed sleep timing, reduced morning alertness, and melatonin suppression.^25^ Importantly, these sex-specific effects were strongly modulated by daytime light intensity: brighter daytime light largely protected females from phase shifts and amplitude dampening, while males were less sensitive to daytime light, maintaining reduced amplitude but without phase alterations. Together, these findings indicate that female sleep-wake regulation is particularly sensitive to the combined effects of nighttime and daytime light, whereas males respond more uniformly with amplitude reduction. Acute DLaN rapidly perturbs sleep continuity in both sexes, while chronic exposure seems to engage circadian mechanisms that reshape sleep-wake organization in a sex- and light-intensity-dependent manner. This unified framework explains why sex differences in sleep-wake behavior are inconsistently observed across rodent studies and highlights daytime light as a critical contextual factor for circadian resilience and vulnerability.

### Sleep pressure under acute and chronic exposure

Acute and chronic DLaN exert different effects on EEG SWA with pronounced sex differences. A single night of DLaN did not produce robust time-dependent changes in NREM sleep SWA in either sex, indicating that acute nighttime light exposure is insufficient to reorganize the temporal structure of SWA. Although modest main effect of DLaN on SWA were observed in both male and female mice, these effects were absent under brighter daytime light, suggesting that sufficient daytime illumination buffers against acute DLaN-induced increased sleep pressure. Overall, acute DLaN data indicates that SWA is relatively resilient to short-term nighttime light exposure, particularly under brighter daytime light. In contrast, chronic DLaN revealed pronounced sex differences in slow-wave sleep regulation. After two weeks of DLaN exposure, males exhibited limited alterations in SWA timing that were restricted to moderate daytime light, whereas females showed robust, time-dependent changes in SWA across all daytime light intensities, particularly during the light phase under dim daytime conditions. Because rapid changes in suprachiasmatic nucleus (SCN) neuronal firing are temporarily correlated with SWA,^26^ these findings suggest that chronic DLaN may alter circadian output signals that regulate sleep, particularly in females.

Human laboratory and real word studies provide important context. Acute exposure to dim light at night reduces slow-wave sleep and increases arousal, and lighter sleep stages in healthy volunteers.^27^ Similar disruptions occur in night-shift workers who frequently experience poor sleep quality and difficulty maintaining consolidated daytime sleep due to circadian misalignment.^28^ Nighttime light exposure during sleep and shift work are both associated with reductions in SWA, a hallmark of deep, restorative sleep, has been linked to impairments in learning and memory ^29^ and increased risk of metabolic disease.^30^ Together, our findings identify SWA as a sensitive marker of nighttime light exposure, particularly in females, and highlight daytime light intensity as a critical factor shaping vulnerability to sleep disruption in modern lighting environments.

### Light intensity matters both day and night: Brighter day and dimmer night

The question whether brighter daytime light can protect sleep under DLaN conditions goes beyond academic interest. DLaN has been shown to induce maladaptive behavioral and physiological outcomes, including reduced sleep efficiency, depressive- and anxiety-like behaviors, metabolic dysfunction and cognitive impairment.^31–33^ Large-scale human studies further demonstrate that exposure to bright nights combined with insufficient daytime light is associated with circadian disruption and increased health risks, including elevated mortality.^9,13,34^ Conversely, greater contrast between bright days and dark nights is associated with earlier and more stable sleep timing.^19^ Together, these observations suggest that sufficient daytime light exposure may buffer the negative consequences of nighttime light.

Our findings provide direct experimental support for this idea. Brighter daytime light attenuated the acute sleep-wake disruption induced by first night DLaN in both sexes, with particularly strong protection in females. Under chronic DLaN (10 lux) conditions, male mice exhibited increased total NREM sleep and reduced wakefulness across all daytime lighting conditions, a result that contrasts with previous studies reporting minimal effects of 5 lux DLaN in mice and rats.^4,16,24,35^ This discrepancy likely reflects the slightly higher nighttime light intensity used here. Consistent with this interpretation, our prior work showed that male and female wildtype mice exposed to 5 lux DLaN under 300 lux daytime lighting did not exhibit significant changes in sleep after six weeks of exposure.^4^

Together, these findings suggest a threshold-like relationship between nighttime light intensity and sleep-wake regulation: very low levels of nighttime illumination may be insufficient to disrupt overall sleep-wake balance, whereas modestly higher levels are sufficient to alter sleep architecture. More broadly, our results support the concept that brighter days combined with darker nights promote sleep and circadian health.

### Mechanistic considerations

The mechanisms underlying this sex-specific circadian vulnerability to light at night are likely multifactorial and may arise at several levels of the circadian system. One possibility is differential sensitivity to photic input pathways, including sex-dependent modulation of retinal signaling, melanopsin-expressing intrinsically photosensitive retinal ganglion cells, or downstream processing within the SCN. Alternatively, sex differences may emerge downstream of the core molecular clock, at the level of circadian output pathways that regulate sleep-wake timing, slow-wave activity, and arousal thresholds. Our observation that males primarily exhibit dampened circadian amplitude whereas females show pronounced phase advances supports the idea that DLaN engages distinct regulatory nodes in the two sexes. The marked reorganization of SWA timing in females further suggests altered coupling between circadian phase and sleep homeostasis, rather than a simple change in sleep pressure. Importantly, the protective effect of brighter daytime light argues that enhanced daytime photic drive may stabilize circadian output signals, thereby buffering against nocturnal light-induced disruption, particularly in females.

### Limitations

Several limitations of this study should be considered. First, we focused on C57BL/6J mice under daytime light intensities commonly used in laboratory settings, circadian and sleep responses to light can vary across strains, and C57BL/6J mice do not synthesize melatonin. Given that light at night can suppress melatonin secretion, future studies should extend these findings to additional strains and species, including melatonin-proficient rodents (e.g., rats) and diurnal species. Second, we did not explicitly control for the estrous cycle stage which could contribute to variability in female responses. Because females spend approximately 20–25% of time in proestrus, EEG recordings obtained without estrous staging are more likely to reflect non-proestrus phases. The robustness and consistency of the observed finding across individuals suggest that estrous-related variability alone is unlikely to account for the magnitude of the effects. Incorporating estrous cycle monitoring in future studies will be important for more accurately assessing sex-specific responses to daytime light intensity and DLaN. Third, our study was conducted under stable light-dark conditions, and different results may emerge under conditions of circadian misalignment, jet lag, or constant light. Finally, while our data implicate circadian reorganization as a key mechanism underlying chronic DLaN effects, direct measurements of SCN molecular rhythms and downstream neural circuits will be required to define the precise loci of sex-specific vulnerability.

## Conclusion

Together, these findings identify daytime light intensity as a critical but underappreciated determinant of sex differences in sleep-wake regulation and vulnerability to light at night. By systematically varying daytime illumination within the range commonly used in laboratory environments, we demonstrate that sex differences in sleep-wake behavior are not fixed properties but emerge from interactions between biological sex, circadian regulation, and environmental light context. Brighter daytime light not only minimizes baseline sex differences but also protects against both acute and chronic DLaN-induced sleep disruption, particularly in females. These results provide a unifying framework to reconcile inconsistencies in the rodent sleep literatures and highlight the importance of reporting and controlling daytime light intensity in circadian and sleep studies. More broadly our findings underscore the potential for daytime light exposure to shape resilience or vulnerability to nocturnal light pollution, with implications for understanding sex-specific sleep and circadian disturbances in modern light environments.

## Resource availability

### Lead contact

Requests for further information and resources should be directed to and will be fulfilled by the lead contact, Christopher S. Colwell.

## Data availability

The datasets here used and analyzed are available from the corresponding author upon request.

## Author contributions

Conceptualization, Y.W., T.D. and C.S.C.; funding acquisition, K.N.P. and G.D.B.; experiment conduction, Y.W. and C.T.C.; data analyzing: Y.W. and T.D.; writing – original draft, Y.W., writing – review and editing, all authors.

## Acknowledgements

We are grateful to Dr. Joanne Zahorsky-Reeves, the veterinarian from the animal facility, for providing post-surgical care for the mice. This study is supported by UCLA Research Support Fund to GDB.

## Ethical approval

All animal procedures were performed in accordance with the UCLA animal care committee’s regulations.

## References

1. Dib, R., Gervais, N.J., and Mongrain, V. (2021). A review of the current state of knowledge on sex differences in sleep and circadian phenotypes in rodents. Neurobiol. Sleep Circadian Rhythms 11, 100068. 10.1016/j.nbscr.2021.100068.

2. Blxler, E.O., Papallaga, M.N., Vgontzas, A.N., Lin, H.-M., Pejovic, S., Kartaraki, M., Vela-Bueno, A., and Chrousos, G.P. (2009). Women sleep objectively better than men and the sleep of young women is more resilient to external stressors: effects of age and menopause. J. Sleep Res. 18, 221–228. 10.1111/j.1365-2869.2008.00713.x.

3. Ehlen, J.C., Hesse, S., Pinckney, L., and Paul, K.N. (2013). Sex Chromosomes Regulate Nighttime Sleep Propensity during Recovery from Sleep Loss in Mice. PLoS ONE 8, e62205. 10.1371/journal.pone.0062205.

4. Wang, Y., Paul, K.N., Block, G.D., Deboer, T., and Colwell, C.S. (2025). Dim light at night disrupts the sleep-wake cycle and exacerbates abnormal EEG activity in Cntnap2 knockout mice: implications for autism spectrum disorders. Mol. Autism 16, 62. 10.1186/s13229-025-00689-7.

5. Choi, J., Kim, S.J., Fujiyama, T., Miyoshi, C., Park, M., Suzuki-Abe, H., Yanagisawa, M., and Funato, H. (2021). The Role of Reproductive Hormones in Sex Differences in Sleep Homeostasis and Arousal Response in Mice. Front. Neurosci. 15, 739236. 10.3389/fnins.2021.739236.

6. Koehl, M., Battle, S., and Meerlo, P. (2006). Sex Differences in Sleep: the Response to Sleep Deprivation and Restraint Stress in Mice. Sleep 29, 1224–1231. 10.1093/sleep/29.9.1224.

7. Paul, K.N., Dugovic, C., Turek, F.W., and Laposky, A.D. (2006). Diurnal Sex Differences in the Sleep-Wake Cycle of Mice are Dependent on Gonadal Function. Sleep 29, 1211–1223. 10.1093/sleep/29.9.1211.

8. Deng, Q., Li, Y., Sun, Z., Fenqin, X., Gao, X., and Li, R. (2025). Effect of estrogens on sex-specific influence in sleep deprivation in mice. Brain Res. Bull. 233, 111663. 10.1016/j.brainresbull.2025.111663.

9. Windred, D.P., Burns, A.C., Lane, J.M., Olivier, P., Rutter, M.K., Saxena, R., Phillips, A.J.K., and Cain, S.W. (2024). Brighter nights and darker days predict higher mortality risk: A prospective analysis of personal light exposure in >88,000 individuals. Proc. Natl. Acad. Sci. 121, e2405924121. 10.1073/pnas.2405924121.

10. Bano-Otalora, B., Martial, F., Harding, C., Bechtold, D.A., Allen, A.E., Brown, T.M., Belle, M.D.C., and Lucas, R.J. (2021). Bright daytime light enhances circadian amplitude in a diurnal mammal. Proc. Natl. Acad. Sci. 118, e2100094118. 10.1073/pnas.2100094118.

11. Borbély, A.A., Daan, S., Wirz-Justice, A., and Deboer, T. (2016). The two-process model of sleep regulation: a reappraisal. J. Sleep Res. 25, 131–143. 10.1111/jsr.12371.

12. Schweizer, A., Berchtold, A., Barrense-Dias, Y., Akre, C., and Suris, J.-C. (2017). Adolescents with a smartphone sleep less than their peers. Eur. J. Pediatr. 176, 131–136. 10.1007/s00431-016-2823-6.

13. Burns, A.C., Windred, D.P., Rutter, M.K., Olivier, P., Vetter, C., Saxena, R., Lane, J.M., Phillips, A.J.K., and Cain, S.W. (2023). Day and night light exposure are associated with psychiatric disorders: an objective light study in >85,000 people. Nat. Ment. Health 1, 853–862. 10.1038/s44220-023-00135-8.

14. Tam, S.K.E., Brown, L.A., Wilson, T.S., Tir, S., Fisk, A.S., Pothecary, C.A., Van Der Vinne, V., Foster, R.G., Vyazovskiy, V.V., Bannerman, D.M., et al. (2021). Dim light in the evening causes coordinated realignment of circadian rhythms, sleep, and short-term memory. Proc. Natl. Acad. Sci. 118, e2101591118. 10.1073/pnas.2101591118.

15. Wang, H.B., Tahara, Y., Luk, S.H.C., Kim, Y.-S., Hitchcock, O.N., Kaswan, Z.A.M., Kim, Y.I., Block, G.D., Ghiani, C.A., Loh, D.H., et al. (2020). Melatonin treatment of repetitive behavioral deficits in the Cntnap2 mouse model of autism spectrum disorder. Neurobiol. Dis. 145, 105064. 10.1016/j.nbd.2020.105064.

16. Stenvers, D.J., Van Dorp, R., Foppen, E., Mendoza, J., Opperhuizen, A.-L., Fliers, E., Bisschop, P.H., Meijer, J.H., Kalsbeek, A., and Deboer, T. (2016). Dim light at night disturbs the daily sleep-wake cycle in the rat. Sci. Rep. 6, 35662. 10.1038/srep35662.

17. Hughes, M.E., Hogenesch, J.B., and Kornacker, K. (2010). JTK_CYCLE: An Efficient Nonparametric Algorithm for Detecting Rhythmic Components in Genome-Scale Data Sets. J. Biol. Rhythms 25, 372–380. 10.1177/0748730410379711.

18. Brooks, T.G., Mrčela, A., Lahens, N.F., Paschos, G.K., Grosser, T., Skarke, C., FitzGerald, G.A., and Grant, G.R. (2022). Nitecap: An Exploratory Circadian Analysis Web Application. J. Biol. Rhythms 37, 43–52. 10.1177/07487304211054408.

19. Blume, C., Garbazza, C., and Spitschan, M. (2019). Effects of light on human circadian rhythms, sleep and mood. Somnologie 23, 147–156. 10.1007/s11818-019-00215-x.

20. Weiße, I., Stötzer, H., and Seitz, R. (1974). Age- and light-dependent changes in the rat eye. Virchows Arch. A 362, 145–156. 10.1007/BF00432392.

21. Rapp, L.M., and Williams, T.P. (1980). A Parametric Study of Retinal Light Damage in Albino and Pigmented Rats. In The Effects of Constant Light on Visual Processes, T. P. Williams and B. N. Baker, eds. (Springer US), pp. 135–159. 10.1007/978-1-4684-7257-8_6.

22. Cho, C.-H., Lee, H.-J., Yoon, H.-K., Kang, S.-G., Bok, K.-N., Jung, K.-Y., Kim, L., and Lee, E.-I. (2016). Exposure to dim artificial light at night increases REM sleep and awakenings in humans. Chronobiol. Int. 33, 117–123. 10.3109/07420528.2015.1108980.

23. Cho, C.-H., Yoon, H.-K., Kang, S.-G., Kim, L., Lee, E.-I., and Lee, H.-J. (2018). Impact of Exposure to Dim Light at Night on Sleep in Female and Comparison with Male Subjects. Psychiatry Investig. 15, 520–530. 10.30773/pi.2018.03.17.

24. Panagiotou, M., Rohling, J.H.T., and Deboer, T. (2020). Sleep Network Deterioration as a Function of Dim-Light-At-Night Exposure Duration in a Mouse Model. Clocks Sleep 2, 308–324. 10.3390/clockssleep2030023.

25. Chang, A.-M., Aeschbach, D., Duffy, J.F., and Czeisler, C.A. (2015). Evening use of light-emitting eReaders negatively affects sleep, circadian timing, and next-morning alertness. Proc. Natl. Acad. Sci. 112, 1232–1237. 10.1073/pnas.1418490112.

26. Deboer, T., Vansteensel, M.J., Détári, L., and Meijer, J.H. (2003). Sleep states alter activity of suprachiasmatic nucleus neurons. Nat. Neurosci. 6, 1086–1090. 10.1038/nn1122.

27. Cho, J.R., Joo, E.Y., Koo, D.L., and Hong, S.B. (2013). Let there be no light: the effect of bedside light on sleep quality and background electroencephalographic rhythms. Sleep Med. 14, 1422–1425. 10.1016/j.sleep.2013.09.007.

28. Alfonsi, V., Scarpelli, S., Gorgoni, M., Pazzaglia, M., Giannini, A.M., and De Gennaro, L. (2021). Sleep-Related Problems in Night Shift Nurses: Towards an Individualized Interventional Practice. Front. Hum. Neurosci. 15, 644570. 10.3389/fnhum.2021.644570.

29. Walker, M.P. (2009). The Role of Slow Wave Sleep in Memory Processing. J. Clin. Sleep Med. JCSM Off. Publ. Am. Acad. Sleep Med. 5, S20–S26.

30. Tasali, E., Leproult, R., Ehrmann, D.A., and Van Cauter, E. (2008). Slow-wave sleep and the risk of type 2 diabetes in humans. Proc. Natl. Acad. Sci. 105, 1044–1049. 10.1073/pnas.0706446105.

31. Wyse, C.A., Biello, S.M., and Gill, J.M.R. (2014). The bright-nights and dim-days of the urban photoperiod: implications for circadian rhythmicity, metabolism and obesity. Ann. Med. 46, 253–263. 10.3109/07853890.2014.913422.

32. Cissé, Y.M., Peng, J., and Nelson, R.J. (2016). Dim Light at Night Prior to Adolescence Increases Adult Anxiety-like Behaviors. Chronobiol. Int. 33, 1473–1480. 10.1080/07420528.2016.1221418.

33. Yook, S., Choi, S.J., Lee, H., Joo, E.Y., and Kim, H. (2024). Long-term night-shift work is associated with accelerates brain aging and worsens N3 sleep in female nurses. Sleep Med. 121, 69–76. 10.1016/j.sleep.2024.06.013.

34. Xiao, Q., Gee, G., Jones, R.R., Jia, P., James, P., and Hale, L. (2020). Cross-sectional association between outdoor artificial light at night and sleep duration in middle-to-older aged adults: The NIH-AARP Diet and Health Study. Environ. Res. 180, 108823. 10.1016/j.envres.2019.108823.

35. Borniger, J.C., Weil, Z.M., Zhang, N., and Nelson, R.J. (2013). Dim Light at Night Does Not Disrupt Timing or Quality of Sleep in Mice. Chronobiol. Int. 30, 1016–1023. 10.3109/07420528.2013.803196.

