## Supplemental figures 1-5 and tables 1-6 for "Bright Days Buffer Nighttime Light: Daytime Illumination Shapes Sex Differences in Sleep and Circadian Regulation"

### Supplemental figures1-5

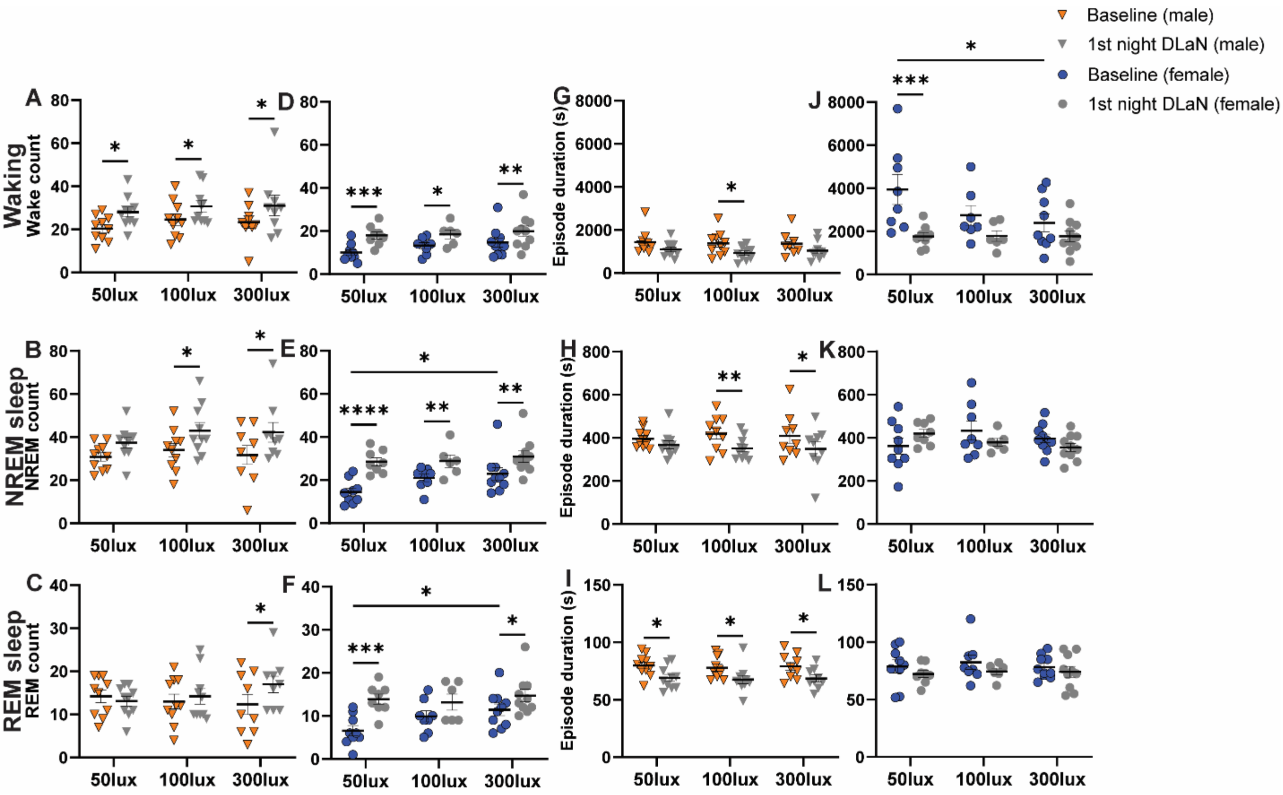


**Figure S1. Bout number there are notre are notre are not duration in male and female WT mice during the first night of DLaN under different daytime light intensities.**

(**A–F**) Number of bouts of wakefulness, NREM sleep, and REM sleep during the night in male (**A-C**) and female (**D-F**) mice exposed to 50, 100, or 300 lux daytime light. (**G-L**) Episode duration of wakefulness, NREM sleep, and REM sleep during the night in male (**G-I**) and female (**J-L**) mice exposed to 50, 100, or 300 lux daytime light. Asterisks indicate significant differences across daytime light intensity or 1st night DLaN (Bonferroni multiple-comparisons test after two-way RM ANOVA or Mixed-effects model; * $p < 0.05$, ** $p < 0.01$, *** $p < 0.001$, *** $p < 0.0001$). Data are presented as mean ± SEM. Sample sizes: 50 lux, 10 males and 9 females; 100 lux, 10 males and 8 females (first-night DLaN: 7 females); 300 lux, 10 males and 10 females (first-night DLaN: 9 males and 9 females). Data are presented as mean ± SEM.

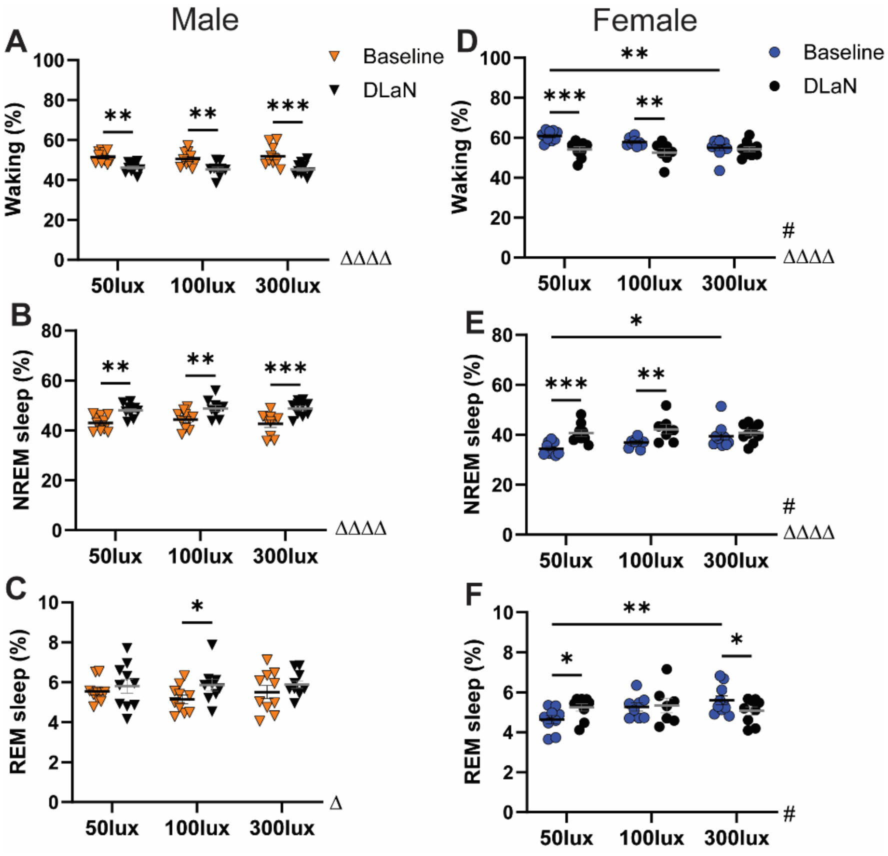


**Figure S2. Total amount of wakefulness and sleep in male and female WT mice during baseline and DLaN under different daytime light intensities.**

(**A-C**) Total amount of wakefulness (**A**), NREM sleep (**B**), and REM sleep (**C**) in male mice under baseline conditions (orange triangles) and after 2 weeks of DLaN exposure (black triangles) following exposure to 50, 100, or 300 lux daytime light. (**D-F**) Total amount of wakefulness (**D**), NREM sleep (**E**), and REM sleep (**F**) in female mice under baseline conditions (blue circles) and after 2 weeks of DLaN exposure (black circles) following exposure to 50, 100, or 300 lux daytime light. Pound signs (#) indicate a significant interaction between daytime light intensity and DLaN (Mixed-effects model with Geisser–Greenhouse correction; # $p < 0.05$). Main effects of DLaN is indicated by Δ (Δ $p < 0.05$, ΔΔΔΔ $p < 0.0001$). Asterisks denote significant post hoc differences between baseline and DLaN (Bonferroni multiple-comparisons test; * $p < 0.05$, ** $p < 0.01$, *** $p < 0.001$).

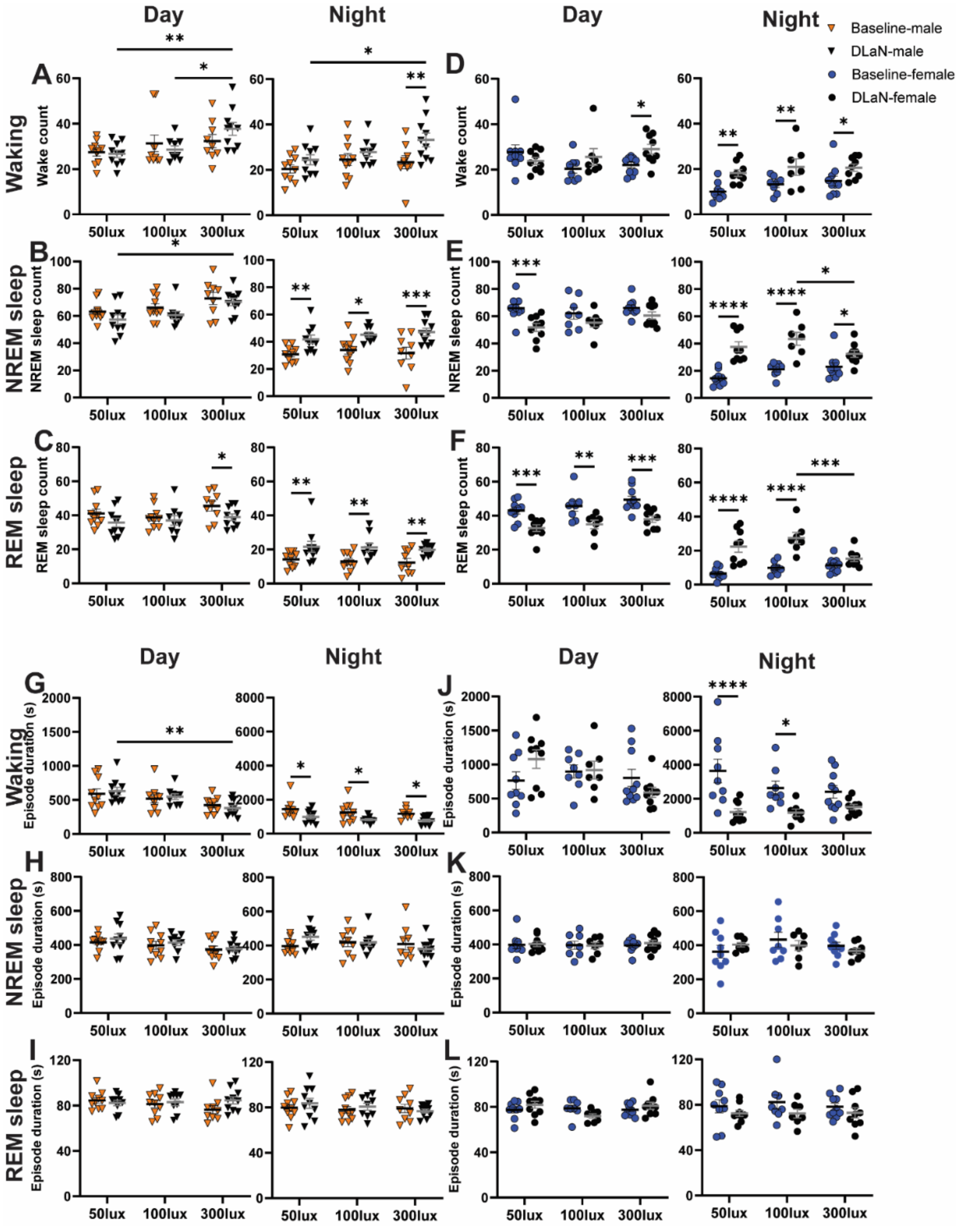


**Figure S3. Bout number and duration in male and female WT mice during 2 weeks of DLaN under different daytime light intensities.**

(**A–F**) Number of bouts of wakefulness, NREM sleep, and REM sleep during the day (left) and the night (right) in male (**A-C**) and female (**D-F**) mice exposed to 50, 100, or 300 lux daytime light. (**G-L**) Episode duration of wakefulness, NREM sleep, and REM sleep during the day (left) and the night (right) in male (**G-I**) and female (**J-L**) mice exposed to 50, 100, or 300 lux daytime light. Asterisks indicate significant differences across daytime light intensity or DLaN (Bonferroni multiple-comparisons test after Mixed-effects model; * $p < 0.05$, ** $p < 0.01$, *** $p < 0.001$, *** $p < 0.0001$). Data are presented as mean ± SEM. Sample sizes: 50 lux, 10 males and 9 females; 100 lux, 10 males and 8 females (DLaN: 7 females); 300 lux, 10 males and 10 females (DLaN: 9 males and 9 females). Data are presented as mean ± SEM.

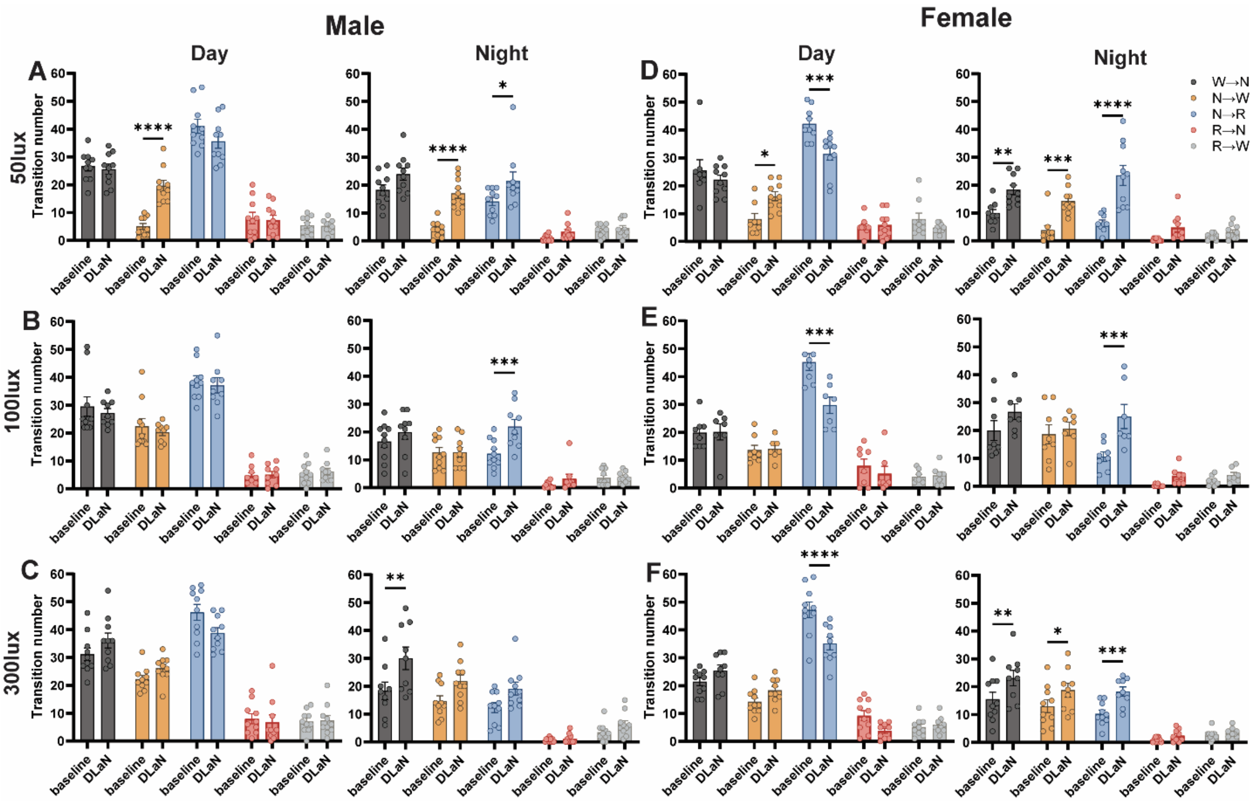


**Figure S4. State transitions in male and female WT mice during 2 weeks of DLaN under different daytime light intensities.**

(**A–F**) Number of bouts of state transitions during the day (left) and the night (right) in male (**A-C**) and female (**D-F**) mice exposed to 50, 100, or 300 lux daytime light. Asterisks indicate significant differences between baseline and DLaN (Bonferroni multiple-comparisons test after Mixed-effects model; * $p < 0.05$, ** $p < 0.01$, *** $p < 0.001$, *** $p < 0.0001$). Data are presented as mean ± SEM. Sample sizes: 50 lux, 10 males and 9 females; 100 lux, 10 males and 8 females (DLaN: 7 females); 300 lux, 10 males and 10 females (DLaN: 9 males and 9 females). Data are presented as mean ± SEM.

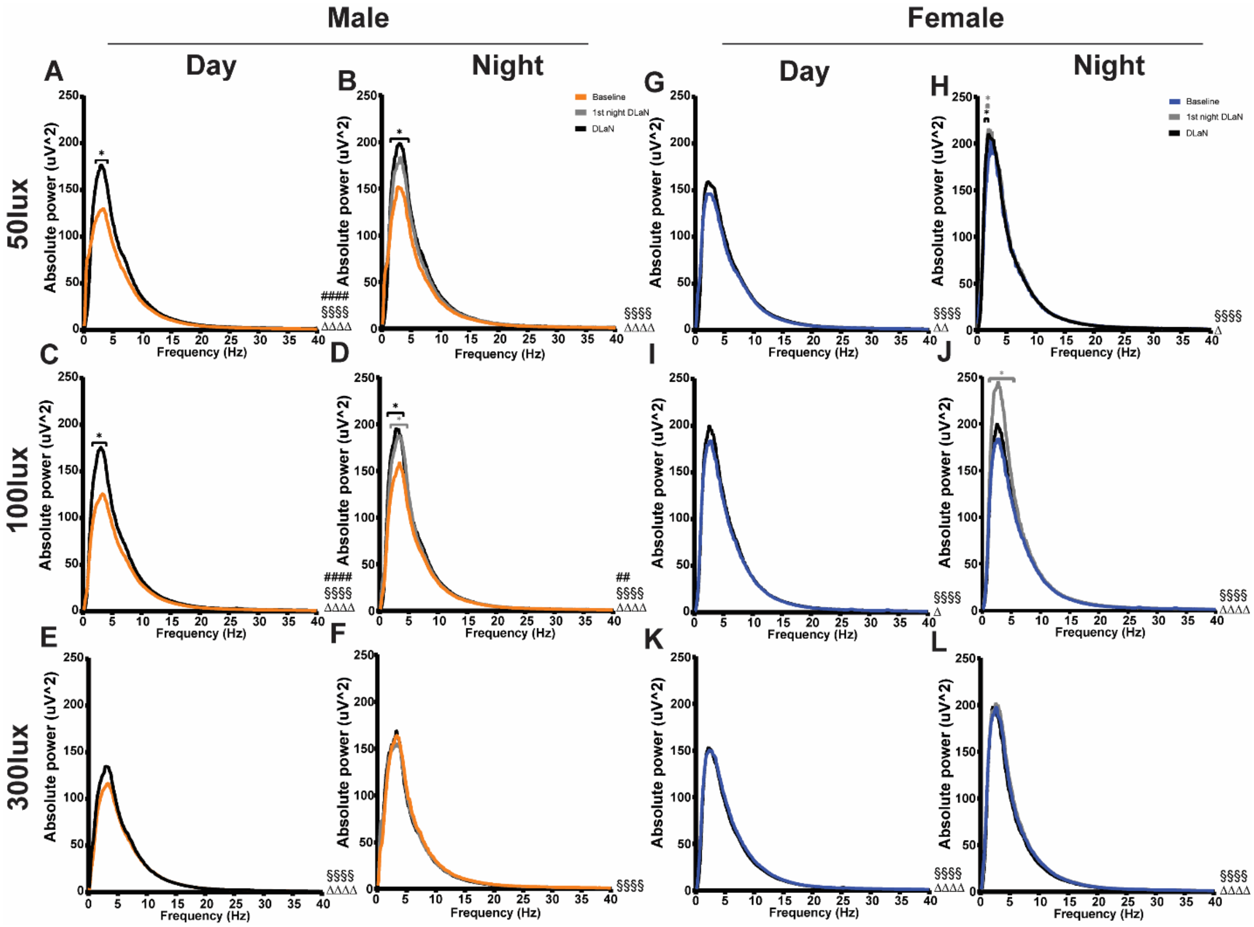


**Figure S5. Absolute power spectrum of NREM sleep in male and female WT mice exposed to DLaN under different daytime light intensities.**

Absolute NREM sleep power spectrum during the daytime are shown for male (**A, C, E**) and female (**G, I, K**) mice following exposure to 50, 100, or 300 lux daytime light under baseline conditions (orange) and after 2 weeks of DLaN exposure (black). Absolute NREM sleep power spectrum during the nighttime are shown for male (**B, D, F**) and female (**H, J, L**) mice following exposure to 50, 100, or 300 lux daytime light under baseline conditions (orange), during the first night of DLaN exposure (gray), and after 2 weeks of DLaN exposure (black). Pound signs (#) indicate a significant interaction between Frequency and DLaN (two-way ANOVA; ## $p < 0.01$, ### $p < 0.001$). Main effects of Frequency and DLaN are indicated by § and Δ, respectively (§§§§ $p < 0.0001$; Δ $p < 0.05$, ΔΔ $p < 0.01$, ΔΔΔΔ, $p < 0.0001$). Grey asterisks denote significant post hoc differences between baseline and 1st night DLaN conditions, black asterisks denote significant post hoc differences between baseline and DLaN conditions (Bonferroni multiple-comparisons test; * $p < 0.05$).

### Supplemental tables 1-6

**Table 1. Statistical difference in Figure 1 data**

| Description | Vigilance state/day night | Statistical test | Sample size | Statistical results | P value |
|---|---|---|---|---|---|
| 50lux daylight | Waking | RM two way ANOVA | 10 male and 9 female | Interaction: F (23, 391) = 2.236 | ***p* =0.0010** |
| | | | | Time effect: F (23, 391) = 32.24 | ***p* <0.0001** |
| | | | | Sex effect: F (1, 17) = 50.38 | ***p* <0.0001** |
| | NREM sleep | RM two way ANOVA | | Interaction: F (23, 391) = 2.339 | ***p* =0.0005** |
| | | | | Time effect: F (23, 391) = 30.93 | ***p* <0.0001** |
| | | | | Sex effect: F (1, 17) = 41.13 | ***p* <0.0001** |
| | REM sleep | RM two way ANOVA | | Interaction: F (23, 391) = 1.373 | *p* =0.1188 |
| | | | | Time effect: F (8.023, 136.4) = 24.70 | ***p* <0.0001** |
| | | | | Sex effect: F (1, 17) = 11.32 | ***p* =0.0037** |
| 100lux daylight | Waking | RM two way ANOVA | 10 male and 8 female | Interaction: F (23, 368) = 1.056 | *p* =0.3933 |
| | | | | Time effect: F (8.942, 143.1) = 19.85 | ***p* <0.0001** |
| | | | | Sex effect: F (1, 16) = 25.58 | ***p* =0.0001** |
| | NREM sleep | RM two way ANOVA | | Interaction: F (23, 368) = 1.010 | *p* =0.4518 |
| | | | | Time effect: F (9.054, 144.9) = 18.25 | ***p* <0.0001** |
| | | | | Sex effect: F (1, 16) = 27.85 | ***p* <0.0001** |
| | REM sleep | RM two way ANOVA | | Interaction: F (23, 368) = 1.291 | *p* =0.1687 |
| | | | | Time effect: F (8.054, 128.9) = 20.59 | ***p* <0.0001** |
| | | | | Sex effect: F (1, 16) = 0.1580 | *p* =0.6962 |
| 300lux daylight | Waking | RM two way ANOVA | 10 male and 10 female | Interaction: F (23, 414) = 1.190 | *p* =0.2488 |
| | | | | Time effect: F (8.548, 153.9) = 29.78 | ***p* <0.0001** |
| | | | | Sex effect: F (1, 18) = 2.048 | *p* =0.1696 |
| | NREM sleep | RM two way ANOVA | | Interaction: F (23, 414) = 1.171 | *p* =0.2667 |
| | | | | Time effect: F (8.485, 152.7) = 27.70 | ***p* <0.0001** |
| | | | | Sex effect: F (1, 18) = 2.421 | *p* =0.1371 |
| | REM sleep | RM two way ANOVA | | Interaction: F (23, 414) = 0.9808 | *p* =0.4893 |
| | | | | Time effect: F (8.590, 154.6) = 25.16 | ***p* <0.0001** |
| | | | | Sex effect: F (1, 18) = 0.06111 | *p* =0.8075 |
| Waking | Day | Two way ANOVA | 50lux: | Interaction: F (2, 51) = 0.1322 | *p* =0.8764 |
| | | | 10 male and 9 female | light intensity effect: F (2, 51) = 2.903 | *p* =0.0639 |
| | | | 100lux: | Sex effect: F (1, 51) = 1.829 | *p* =0.1822 |
| | Night | Two way ANOVA | 10 male and 8 female | Interaction: F (2, 51) = 3.465 | ***p* =0.0388** |
| | | | 300lux: | light intensity effect: F (2, 51) = 3.637 | ***p* =0.0334** |
| | | | 10 male and 10 female | Sex effect: F (1, 51) = 40.10 | ***p* <0.0001** |
| NREM sleep | Day | Two way ANOVA | 50lux: | Interaction: F (2, 51) = 0.3876 | *p* =0.6807 |
| | | | 10 male and 9 female | light intensity effect: F (2, 51) = 2.433 | *p* =0.0979 |
| | | | 100lux: | Sex effect: F (1, 51) = 3.468 | *p* =0.0683 |
| | Night | Two way ANOVA | 10 male and 8 female | Interaction: F (2, 51) = 3.465 | ***p* =0.0388** |
| | | | 300lux: | light intensity effect: F (2, 51) = 3.637 | ***p* =0.0334** |
| | | | 10 male and 10 female | Sex effect: F (1, 51) = 40.10 | ***p* <0.0001** |
| REM Sleep | Day | Two way ANOVA | 50lux: | Interaction: F (2, 51) = 0.7793 | *p* =0.4641 |
| | | | 10 male and 9 female | light intensity effect: F (2, 51) = 1.641 | *p* =0.2038 |
| | | | 100lux: | Sex effect: F (1, 51) = 0.9380 | *p* =0.3374 |
| | Night | Two way ANOVA | 10 male and 8 female | Interaction: F (2, 51) = 4.026 | ***p* =0.0238** |
| | | | 300lux: | light intensity effect: F (2, 51) = 0.5240 | *p* =0.5953 |
| | | | 10 male and 10 female | Sex effect: F (1, 51) = 11.78 | ***p* =0.0012** |

**Table 2. Statistical difference in Figure 2 data**

| Description | Vigilance state/day night | Statistical test | Sample size | Statistical results | P value |
|---|---|---|---|---|---|
| Bout numbers of Waking | Day | Two way ANOVA | 50lux: | Interaction: F (2, 50) = 2.927 | $p$ =0.0628 |
| | | | 10 male and 9 female | light intensity effect: F (2, 50) = 0.2283 | $p$ =0.7967 |
| | | | 100lux: | Sex effect: F (1, 50) = 10.74 | **$p$ =0.0019** |
| | Night | Two way ANOVA | 10 male and 8 female | Interaction: F (2, 50) = 0.1897 | $p$ =0.8278 |
| | | | 300lux: | light intensity effect: F (2, 50) = 2.023 | $p$ =0.1430 |
| | | | 9 male and 10 female | Sex effect: F (1, 50) = 32.61 | **$p$ <0.0001** |
| Bout numbers of NREM sleep | Day | Two way ANOVA | 50lux: | Interaction: F (2, 50) = 1.214 | $p$ =0.3055 |
| | | | 10 male and 9 female | light intensity effect: F (2, 50) = 1.784 | $p$ =0.1784 |
| | | | 100lux: | Sex effect: F (1, 50) = 1.033 | $p$ =0.3144 |
| | Night | Two way ANOVA | 10 male and 8 female | Interaction: F (2, 50) = 0.9121 | $p$ =0.4083 |
| | | | 300lux: | light intensity effect: F (2, 50) = 1.8937 | $p$ =0.1612 |
| | | | 9 male and 10 female | Sex effect: F (1, 50) = 29.90 | **$p$ <0.0001** |
| Bout numbers of REM sleep | Day | Two way ANOVA | 50lux: | Interaction: F (2, 50) = 0.5078 | $p$ =0.6049 |
| | | | 10 male and 9 female | light intensity effect: F (2, 50) = 3.180 | $p$ =0.0501 |
| | | | 100lux: | Sex effect: F (1, 50) = 4.509 | **$p$ =0.0387** |
| | Night | Two way ANOVA | 10 male and 8 female | Interaction: F (2, 50) = 2.321 | $p$ =0.1087 |
| | | | 300lux: | light intensity effect: F (2, 50) = 0.4653 | $p$ =0.6307 |
| | | | 9 male and 10 female | Sex effect: F (1, 50) = 8.868 | **$p$ =0.0045** |
| Episode duration of Waking | Day | Two way ANOVA | 50lux: | Interaction: F (2, 50) = 0.8099 | $p$ =0.4507 |
| | | | 10 male and 9 female | light intensity effect: F (2, 50) = 0.5139 | $p$ =0.6013 |
| | | | 100lux: | Sex effect: F (1, 50) = 16.52 | **$p$ =0.0002** |
| | Night | Two way ANOVA | 10 male and 8 female | Interaction: F (2, 49) = 0.9768 | $p$ =0.3837 |
| | | | 300lux: | light intensity effect: F (2, 49) = 2.276 | $p$ =0.1134 |
| | | | 9 male and 10 female | Sex effect: F (1, 49) = 27.40 | **$p$ <0.0001** |
| Episode duration of NREM sleep | Day | Two way ANOVA | 50lux: | Interaction: F (2, 50) = 0.5471 | $p$ =0.5821 |
| | | | 10 male and 9 female | light intensity effect: F (2, 50) = 0.7561 | $p$ =0.4748 |
| | | | 100lux: | Sex effect: F (1, 50) = 0.001228 | $p$ =0.9722 |
| | Night | Two way ANOVA | 10 male and 8 female | Interaction: F (2, 50) = 0.3465 | $p$ =0.7089 |
| | | | 300lux: | light intensity effect: F (2, 50) = 1.245 | $p$ =0.2968 |
| | | | 9 male and 10 female | Sex effect: F (1, 50) = 0.1875 | $p$ =0.6668 |
| Episode duration of REM sleep | Day | Two way ANOVA | 50lux: | Interaction: F (2, 50) = 1.250 | $p$ =0.2953 |
| | | | 10 male and 9 female | light intensity effect: F (2, 50) = 1.178 | $p$ =0.3162 |
| | | | 100lux: | Sex effect: F (1, 50) = 1.689 | $p$ =0.1997 |
| | Night | Two way ANOVA | 10 male and 8 female | Interaction: F (2, 50) = 0.2889 | $p$ =0.7504 |
| | | | 300lux: | light intensity effect: F (2, 50) = 0.05977 | $p$ =0.9421 |
| | | | 9 male and 10 female | Sex effect: F (1, 50) = 0.06160 | $p$ =0.8050 |

**Table 3. Statistical difference in Figure 3 data**

| Description | Vigilance state | Statistical test | Sample size | Statistical results | P value |
|---|---|---|---|---|---|
| 50lux daylight (male) | Waking | RM two way ANOVA | n =10 | Interaction: F (23, 207) = 3.473 | ***p* <0.0001** |
| | | | | Time effect: F (23, 207) = 21.53 | ***p* <0.0001** |
| | | | | 1st night DLaN effect: F (1, 9) = 1.017 | *p* =0.3397 |
| | NREM sleep | RM two way ANOVA | | Interaction: F (23, 207) = 3.589 | ***p* <0.0001** |
| | | | | Time effect: F (23, 207) = 19.44 | ***p* <0.0001** |
| | | | | 1st night DLaN effect: F (1, 9) = 2.028 | *p* =0.1882 |
| | REM sleep | RM two way ANOVA | | Interaction: F (23, 207) = 2.294 | *p* =0.1739 |
| | | | | Time effect: F (23, 207) = 19.87 | ***p* =0.0011** |
| | | | | 1st night DLaN effect: F (1, 9) = 2.180 | ***p* <0.0001** |
| 100lux daylight (male) | Waking | RM two way ANOVA | n=10 | Interaction: F (23, 207) = 3.000 | ***p* <0.0001** |
| | | | | Time effect: F (23, 207) = 15.93 | ***p* <0.0001** |
| | | | | 1st night DLaN effect: F (1, 9) = 0.5497 | *p* =0.4773 |
| | NREM sleep | RM two way ANOVA | | Interaction: F (23, 207) = 2.951 | ***p* <0.0001** |
| | | | | Time effect: F (23, 207) = 13.91 | ***p* <0.0001** |
| | | | | 1st night DLaN effect: F (1, 9) = 0.6592 | *p* =0.4378 |
| | REM sleep | RM two way ANOVA | | Interaction: F (6.385, 57.46) = 2.205 | *p* =0.0519 |
| | | | | Time effect: F (5.558, 50.02) = 17.51 | ***p* <0.0001** |
| | | | | 1st night DLaN effect: F (1.000, 9.000) = 0.08425 | *p* =0.7782 |
| 300lux daylight (male) | Waking | Mixed-effects model (REML) | baseline: n=10; 1st night DLaN: n=9 | Interaction: F (23, 183) = 2.044 | ***p* =0.005** |
| | | | | Time effect: F (23, 207) = 22.54 | ***p* <0.0001** |
| | | | | 1st night DLaN effect: F (1, 9) = 3.213 | *p* =0.1067 |
| | NREM sleep | Mixed-effects model (REML) | | Interaction: F (23, 183) = 2.114 | ***p* =0.0034** |
| | | | | Time effect: F (23, 207) = 19.74 | ***p* <0.0001** |
| | | | | 1st night DLaN effect: F (1, 9) = 2.706 | *p* =0.1344 |
| | REM sleep | Mixed-effects model (REML) | | Interaction: F (23, 183) = 1.485 | ***p* =0.0798** |
| | | | | Time effect: F (23, 207) = 23.10 | ***p* <0.0001** |
| | | | | 1st night DLaN effect: F (1, 9) = 3.064 | *p* =0.114 |
| 50lux daylight (female) | Waking | RM two way ANOVA | n =9 | Interaction: F (23, 184) = 2.619 | ***p* =0.0002** |
| | | | | Time effect: F (23, 184) = 15.76 | ***p* <0.0001** |
| | | | | 1st night DLaN effect: F (1, 8) = 18.28 | ***p* =0.0027** |
| | NREM sleep | RM two way ANOVA | | Interaction: F (23, 184) = 2.687 | ***p* =0.0001** |
| | | | | Time effect: F (23, 184) = 14.38 | ***p* <0.0001** |
| | | | | 1st night DLaN effect: F (1, 8) = 17.89 | ***p* =0.0029** |
| | REM sleep | RM two way ANOVA | | Interaction: F (23, 184) = 1.884 | ***p* =0.0116** |
| | | | | Time effect: F (23, 184) = 16.93 | ***p* <0.0001** |
| | | | | 1st night DLaN effect: F (1, 8) = 0.7287 | *p* =0.4181 |
| 100lux daylight (female) | Waking | Mixed-effects model (REML) | baseline: n=8; 1st night DLaN: n=7 | Interaction: F (23, 137) = 2.455 | ***p* =0.0007** |
| | | | | Time effect: F (23, 161) = 11.81 | ***p* <0.0001** |
| | | | | 1st night DLaN effect: F (1, 7) = 1.632 | *p* =0.2422 |
| | NREM sleep | Mixed-effects model (REML) | | Interaction: F (23, 137) = 2.522 | ***p* =0.0005** |
| | | | | Time effect: F (23, 161) = 10.61 | ***p* <0.0001** |
| | | | | 1st night DLaN effect: F (1, 7) = 2.028 | *p* =0.1974 |
| | REM sleep | Mixed-effects model (REML) | | Interaction: F (23, 137) = 1.768 | ***p* =0.024** |
| | | | | Time effect: F (23, 161) = 15.12 | ***p* <0.0001** |
| | | | | 1st night DLaN effect: F (1, 7) = 0.02584 | *p* =0.8768 |
| 300lux daylight (female) | Waking | Mixed-effects model (REML) | baseline: n=10; 1st night DLaN: n=9 | Interaction: F (23, 183) = 4.022 | ***p* <0.0001** |
| | | | | Time effect: F (23, 207) = 15.29 | ***p* <0.0001** |
| | | | | 1st night DLaN effect: F (1, 9) = 0.01259 | *p* =0.9131 |
| | NREM sleep | Mixed-effects model (REML) | | Interaction: F (23, 183) = 3.979 | ***p* <0.0001** |
| | | | | Time effect: F (23, 207) = 14.63 | ***p* <0.0001** |
| | | | | 1st night DLaN effect: F (1, 9) = 0.006574 | *p* =0.9372 |
| | REM sleep | Mixed-effects model (REML) | | Interaction: F (23, 183) = 3.164 | ***p* <0.0001** |
| | | | | Time effect: F (23, 207) = 17.25 | ***p* <0.0001** |
| | | | | 1st night DLaN effect: F (1, 9) = 0.6336 | *p* =0.4465 |
| Waking | Male | Mixed-effects model (REML) | 50lux(Male): n =10 | Interaction: F (2, 26) = 0.3014 | *p* =0.7424 |
| | | | 100lux(Male): n=10 | light intensity effect: F (2, 27) = 1.449 | *p* =0.2524 |
| | | | 300lux(Male): $n_{baseline}$=10; $n_{DLaN}$=9 | 1st night DLaN effect: F (1, 26) = 4.417 | ***p* =0.0454** |
| | Female | Mixed-effects model (REML) | 50lux(Female):n =9 | Interaction: F (2, 22) = 8.246 | ***p* =0.0021** |
| | | | 50lux(Female): $n_{baseline}$=8; $n_{DLaN}$=7 | light intensity effect: F (2, 24) = 1.358 | *p* =0.2762 |
| | | | 300lux(Female): $n_{baseline}$=10; $n_{DLaN}$=9 | 1st night DLaN effect: F (1, 22) = 47.38 | ***p* <0.0001** |
| NREM sleep | Male | Mixed-effects model (REML) | 50lux(Male): n =10 | Interaction: F (2, 26) = 0.1629 | *p* =0.8505 |
| | | | 100lux(Male): n=10 | light intensity effect: F (2, 27) = 1.642 | *p* =0.2123 |
| | | | 300lux(Male): $n_{baseline}$=10; $n_{DLaN}$=9 | 1st night DLaN effect: F (1, 26) = 5.264 | ***p* =0.0301** |
| | Female | Mixed-effects model (REML) | 50lux(Female):n =9 | Interaction: F (2, 22) = 8.670 | ***p* =0.0017** |
| | | | 50lux(Female): $n_{baseline}$=8; $n_{DLaN}$=7 | light intensity effect: F (2, 24) = 1.144 | *p* =0.3354 |
| | | | 300lux(Female): $n_{baseline}$=10; $n_{DLaN}$=9 | 1st night DLaN effect: F (1, 22) = 48.14 | ***p* <0.0001** |
| REM sleep | Male | Mixed-effects model (REML) | 50lux(Male): n =10 | Interaction: F (2, 26) = 4.234 | ***p* =0.0256** |
| | | | 100lux(Male): n=10 | light intensity effect: F (2, 27) = 0.008742 | *p* =0.9913 |
| | | | 300lux(Male): $n_{baseline}$=10; $n_{DLaN}$=9 | 1st night DLaN effect: F (1, 26) = 0.05730 | *p* =0.8127 |
| | Female | Mixed-effects model (REML) | 50lux(Female):n =9 | Interaction: F (2, 22) = 2.286 | *p* =0.1253 |
| | | | 50lux(Female): $n_{baseline}$=8; $n_{DLaN}$=7 | light intensity effect: F (2, 24) = 3.160 | *p* =0.0605 |
| | | | 300lux(Female): $n_{baseline}$=10; $n_{DLaN}$=9 | 1st night DLaN effect: F (1, 22) = 21.10 | ***p* =0.0001** |

**Table 4. Statistical difference in Figure 4 data**

| Description | Vigilance state | Statistical test | Sample size | Statistical results | P value |
|---|---|---|---|---|---|
| 50lux daylight (male) | Waking | RM two way ANOVA | n =10 | Interaction: F (23, 207) = 5.043 | ***p* <0.0001** |
| | | | | Time effect: F (23, 207) = 10.27 | ***p* <0.0001** |
| | | | | DLaN effect: F (1, 9) = 15.30 | ***p* =0.0036** |
| | NREM sleep | RM two way ANOVA | | Interaction: F (23, 207) = 5.037 | ***p* <0.0001** |
| | | | | Time effect: F (23, 207) = 9.452 | ***p* <0.0001** |
| | | | | DLaN effect: F (1, 9) = 15.59 | ***p* =0.0034** |
| | REM sleep | RM two way ANOVA | | Interaction: F (23, 207) = 3.140 | ***p* <0.0001** |
| | | | | Time effect: F (23, 207) = 9.853 | ***p* <0.0001** |
| | | | | DLaN effect: F (1, 9) = 0.5047 | *p* =0.4954 |
| 100lux daylight (male) | Waking | Mixed-effects model (REML) | baseline: n=10; DLaN: n=9 | Interaction: F (23, 183) = 5.061 | ***p* <0.0001** |
| | | | | Time effect: F (23, 207) = 9.660 | ***p* <0.0001** |
| | | | | DLaN effect: F (1, 9) = 7.867 | ***p* =0.0205** |
| | NREM sleep | Mixed-effects model (REML) | | Interaction: F (23, 183) = 4.806 | ***p* <0.0001** |
| | | | | Time effect: F (23, 207) = 8.758 | ***p* <0.0001** |
| | | | | DLaN effect: F (1, 9) = 7.451 | ***p* =0.0232** |
| | REM sleep | Mixed-effects model (REML) | | Interaction: F (23, 183) = 4.065 | ***p* <0.0001** |
| | | | | Time effect: F (23, 207) = 9.639 | ***p* <0.0001** |
| | | | | DLaN effect: F (1, 9) = 3.805 | *p* =0.0829 |
| 300lux daylight (male) | Waking | RM two way ANOVA | n =10 | Interaction: F (23, 207) = 4.391 | ***p* <0.0001** |
| | | | | Time effect: F (23, 207) = 14.72 | ***p* <0.0001** |
| | | | | DLaN effect: F (1, 9) = 10.62 | ***p* =0.0099** |
| | NREM sleep | RM two way ANOVA | | Interaction: F (1, 408) = 0.1547 | *p* =0.6943 |
| | | | | Time effect: F (23, 408) = 2.842 | ***p* <0.0001** |
| | | | | DLaN effect: F (23, 408) = 22.33 | ***p* <0.0001** |
| | REM sleep | RM two way ANOVA | | Interaction: F (23, 207) = 2.706 | ***p* <0.0001** |
| | | | | Time effect: F (23, 207) = 14.29 | ***p* <0.0001** |
| | | | | DLaN effect: F (1, 9) = 1.114 | *p* =0.3188 |
| 50lux daylight (female) | Waking | RM two way ANOVA | n =9 | Interaction: F (23, 184) = 5.458 | ***p* <0.0001** |
| | | | | Time effect: F (23, 184) = 14.28 | ***p* <0.0001** |
| | | | | DLaN effect: F (1, 8) = 41.19 | ***p* =0.0002** |
| | NREM sleep | RM two way ANOVA | | Interaction: F (23, 184) = 5.223 | ***p* <0.0001** |
| | | | | Time effect: F (23, 184) = 13.56 | ***p* <0.0001** |
| | | | | DLaN effect: F (1, 8) = 49.02 | ***p* =0.0001** |
| | REM sleep | RM two way ANOVA | | Interaction: F (23, 184) = 5.657 | ***p* <0.0001** |
| | | | | Time effect: F (23, 184) = 11.98 | ***p* <0.0001** |
| | | | | DLaN effect: F (1, 8) = 3.887 | *p* =0.0841 |
| 100lux daylight (female) | Waking | Mixed-effects model (REML) | baseline: n=8; DLaN: n=7 | Interaction: F (23, 137) = 2.739 | ***p* =0.0002** |
| | | | | Time effect: F (23, 161) = 9.184 | ***p* <0.0001** |
| | | | | DLaN effect: F (1, 7) = 4.620 | *p* =0.0687 |
| | NREM sleep | Mixed-effects model (REML) | | Interaction: F (23, 137) = 2.434 | ***p* =0.0008** |
| | | | | Time effect: F (23, 161) = 8.717 | ***p* <0.0001** |
| | | | | DLaN effect: F (1, 7) = 5.700 | ***p* =0.0483** |
| | REM sleep | Mixed-effects model (REML) | | Interaction: F (23, 144) = 3.759 | ***p* <0.0001** |
| | | | | Time effect: F (23, 168) = 8.357 | ***p* <0.0001** |
| | | | | DLaN effect: F (1, 144) = 0.03954 | *p* =0.8427 |
| 300lux daylight (female) | Waking | Mixed-effects model (REML) | baseline: n=10; DLaN: n=9 | Interaction: F (23, 183) = 2.853 | **p<0.0001** |
| | | | | Time effect: F (23, 207) = 22.42 | **p<0.0001** |
| | | | | DLaN effect: F (1, 9) = 0.1634 | p=0.6955 |
| | NREM sleep | Mixed-effects model (REML) | | Interaction: F (23, 183) = 2.766 | **p<0.0001** |
| | | | | Time effect: F (23, 207) = 21.66 | **p<0.0001** |
| | | | | DLaN effect: F (1, 9) = 0.6509 | p=0.4406 |
| | REM sleep | Mixed-effects model (REML) | | Interaction: F (23, 408) = 2.367 | **p=0.0004** |
| | | | | Time effect: F (23, 408) = 16.82 | **p<0.0001** |
| | | | | DLaN effect: F (1, 408) = 2.605 | p=0.1073 |
| Waking (day) | Male | Mixed-effects model (REML) | 50lux(Male): n =10 | Interaction: F (2, 26) = 1.728 | *p* =0.1974 |
| | | | 100lux(Male): $n_{baseline}$=10; $n_{DLaN}$=9 | light intensity effect: F (2, 27) = 2.374 | *p* =0.1123 |
| | | | 300lux(Male): n =10 | DLaN effect: F (1, 26) = 12.31 | ***p* =0.0017** |
| | Female | Mixed-effects model (REML) | 50lux(Female):n =9 | Interaction: F (2, 26) = 2.072 | *p* =0.1462 |
| | | | 50lux(Female): $n_{baseline}$=8; $n_{DLaN}$=7 | light intensity effect: F (2, 27) = 1.478 | *p* =0.2459 |
| | | | 300lux(Female): $n_{baseline}$=10; $n_{DLaN}$=9 | DLaN effect: F (1, 26) = 11.53 | ***p*= 0.0022** |
| NREM sleep (day) | Male | Mixed-effects model (REML) | 50lux(Male): n =10 | Interaction: F (2, 26) = 2.049 | *p* =0.1492 |
| | | | 100lux(Male): $n_{baseline}$=10; $n_{DLaN}$=9 | light intensity effect: F (2, 26) = 1.359 | *p* =0.2747 |
| | | | 300lux(Male): n =10 | DLaN effect: F (1, 26) = 11.53 | ***p* =0.0022** |
| | Female | Mixed-effects model (REML) | 50lux(Female):n =9 | Interaction: F (2, 46) = 2.750 | *p* =0.0744 |
| | | | 50lux(Female): $n_{baseline}$=8; $n_{DLaN}$=7 | light intensity effect: F (2, 46) = 4.850 | ***p* =0.0123** |
| | | | 300lux(Female): $n_{baseline}$=10; $n_{DLaN}$=9 | DLaN effect: F (1, 46) = 22.46 | ***p* <0.0001** |
| REM sleep (day) | Male | Mixed-effects model (REML) | 50lux(Male): n =10 | Interaction: F (2, 26) = 0.5273 | *p* =0.5964 |
| | | | 100lux(Male): $n_{baseline}$=10; $n_{DLaN}$=9 | light intensity effect: F (2, 27) = 2.053 | *p* =0.1479 |
| | | | 300lux(Male): n =10 | DLaN effect: F (1, 26) = 1.859 | *p* =0.1845 |
| | Female | Mixed-effects model (REML) | 50lux(Female):n =9 | Interaction: F (2, 22) = 0.5033 | *p* =0.6113 |
| | | | 50lux(Female): $n_{baseline}$=8; $n_{DLaN}$=7 | light intensity effect: F (2, 24) = 2.025 | *p* =0.154 |
| | | | 300lux(Female): $n_{baseline}$=10; $n_{DLaN}$=9 | DLaN effect: F (1, 22) = 49.14 | ***p* <0.0001** |
| Waking (night) | Male | Mixed-effects model (REML) | 50lux(Male): n =10 | Interaction: F (2, 53) = 0.2708 | *p* =0.7638 |
| | | | 100lux(Male): $n_{baseline}$=10; $n_{DLaN}$=9 | light intensity effect: F (2, 53) = 2.130 | *p* =0.129 |
| | | | 300lux(Male): n =10 | DLaN effect: F (1, 53) = 67.42 | ***p* <0.0001** |
| | Female | Mixed-effects model (REML) | 50lux(Female):n =9 | Interaction: F (2, 22) = 6.510 | ***p* =0.006** |
| | | | 50lux(Female): $n_{baseline}$=8; $n_{DLaN}$=7 | light intensity effect: F (2, 24) = 1.304 | *p* =0.29 |
| | | | 300lux(Female): $n_{baseline}$=10; $n_{DLaN}$=9 | DLaN effect: F (1, 22) = 54.62 | ***p* <0.0001** |
| NREM sleep (night) | Male | Mixed-effects model (REML) | 50lux(Male): n =10 | Interaction: F (2, 53) = 0.3544 | *p* =0.7032 |
| | | | 100lux(Male): $n_{baseline}$=10; $n_{DLaN}$=9 | light intensity effect: F (2, 53) = 2.214 | *p* =0.1193 |
| | | | 300lux(Male): n =10 | DLaN effect: F (1, 53) = 66.64 | ***p* <0.0001** |
| | Female | Mixed-effects model (REML) | 50lux(Female):n =9 | Interaction: F (2, 22) = 6.352 | ***p* =0.0066** |
| | | | 50lux(Female): $n_{baseline}$=8; $n_{DLaN}$=7 | light intensity effect: F (2, 24) = 1.172 | *p* =0.3267 |
| | | | 300lux(Female): $n_{baseline}$=10; $n_{DLaN}$=9 | DLaN effect: F (1, 22) = 55.11 | ***p* <0.0001** |
| REM sleep (night) | Male | Mixed-effects model (REML) | 50lux(Male): n =10 | Interaction: F (2, 26) = 17.57 | ***p* <0.0001** |
| | | | 100lux(Male): $n_{baseline}$=10; $n_{DLaN}$=9 | light intensity effect: F (2, 27) = 3.482 | ***p* =0.0452** |
| | | | 300lux(Male): n =10 | DLaN effect: F (1, 26) = 25.68 | ***p* <0.0001** |
| | Female | Mixed-effects model (REML) | 50lux(Female):n =9 | Interaction: F (2, 46) = 6.034 | ***p* =0.0047** |
| | | | 50lux(Female): $n_{baseline}$=8; $n_{DLaN}$=7 | light intensity effect: F (2, 46) = 1.427 | *p* =0.2506 |
| | | | 300lux(Female): $n_{baseline}$=10; $n_{DLaN}$=9 | DLaN effect: F (1, 46) = 38.57 | ***p* <0.0001** |

**Table 5. Statistical difference in Figure 5 data**

| Description | Light intensity | Statistical test | Sample size | Statistical results | P value |
|---|---|---|---|---|---|
| Male (1st night DLaN) | 50lux | Mixed-effects model (REML) | $n_{baseline}$= 10 | Interaction: F (2.495, 17.90) = 1.708 | $p$ = 0.2058 |
| | | | $n_{1st\ night\ DLaN}$= 10 | Time effect: F (2.510, 22.59) = 8.243 | **$p$ = 0.0011** |
| | | | | 1st night DLaN effect: F (1.000, 9.000) = 0.9973 | $p$ = 0.344 |
| | 100lux | Mixed-effects model (REML) | $n_{baseline}$= 9 | Interaction: F (3.008, 20.40) = 2.371 | $p$ = 0.1001 |
| | | | $n_{1st\ night\ DLaN}$= 9 | Time effect: F (3.722, 29.78) = 9.324 | **$p$ < 0.0001** |
| | | | | 1st night DLaN effect: F (1.000, 8.000) = 14.97 | **$p$ = 0.0047** |
| | 300lux | Mixed-effects model (REML) | $n_{baseline}$= 8 | Interaction: F (2.255, 12.75) = 0.4752 | $p$ =0.6543 |
| | | | $n_{1st\ night\ DLaN}$= 8 | Time effect: F (1.927, 13.49) = 8.393 | **$p$ = 0.0046** |
| | | | | 1st night DLaN effect: F (1.000, 7.000) = 2.985 | $p$ = 0.1276 |
| Female (1st night DLaN) | 50lux | Mixed-effects model (REML) | $n_{baseline}$= 9 | - | - |
| | | | $n_{1st\ night\ DLaN}$= 9 | Time effect: F (5.351, 70.96) = 13.83 | **$p$ <0.0001** |
| | | | | 1st night DLaN effect: F (1, 16) = 6.649 | **$p$ =0.0202** |
| | 100lux | Mixed-effects model (REML) | $n_{baseline}$= 8 | Interaction: F (23.00, 61.00) = 1.900 | **$p$ = 0.0242** |
| | | | $n_{1st\ night\ DLaN}$= 6 | Time effect: F (23.00, 161.0) = 8.477 | **$p$ < 0.0001** |
| | | | | - | - |
| | 300lux | Mixed-effects model (REML) | $n_{baseline}$= 10 | Interaction: F (2.473, 11.29) = 2.675 | $p$ = 0.1046 |
| | | | $n_{1st\ night\ DLaN}$= 9 | Time effect: F (2.267, 20.40) = 22.31 | **$p$ <0.0001** |
| | | | | 1st night DLaN effect: F (1.000, 9.000) = 1.037 | $p$ = 0.335 |
| Male (DLaN) | 50lux | Mixed-effects model (REML) | $n_{baseline}$= 10 | Interaction: F (3.846, 28.43) = 2.233 | $p$ =0.0926 |
| | | | $n_{DLaN}$= 10 | Time effect: F (3.406, 30.65) = 8.796 | **$p$ <0.0001** |
| | | | | DLaN effect: F (1, 9) = 5.067 | $p$ =0.0509 |
| | 100lux | Mixed-effects model (REML) | $n_{baseline}$= 9 | Interaction: F (23, 133) = 1.744 | **$p$ = 0.0272** |
| | | | $n_{DLaN}$= 8 | Time effect: F (23, 184) = 2.375 | **$p$ = 0.0008** |
| | | | | DLaN effect: F (1, 8) = 6.502 | **$p$ = 0.0342** |
| | 300lux | Mixed-effects model (REML) | $n_{baseline}$= 8 | Interaction: F (2.575, 15.23) = 1.032 | $p$ = 0.3968 |
| | | | $n_{DLaN}$= 8 | Time effect: F (3.093, 21.65) = 8.744 | **$p$ = 0.0005** |
| | | | | DLaN effect: F (1.000, 7.000) = 3.501 | $p$ = 0.1035 |
| Female (DLaN) | 50lux | Mixed-effects model (REML) | $n_{baseline}$= 9 | - | - |
| | | | $n_{DLaN}$= 9 | Time effect: F (23, 313) = 6.238 | **$p$ <0.0001** |
| | | | | DLaN effect: F (2, 16) = 10.24 | **$p$ <0.0001** |
| | 100lux | Mixed-effects model (REML) | $n_{baseline}$= 8 | Interaction: F (23.00, 84.00) = 3.378 | **$p$ = 0.0011** |
| | | | $n_{DLaN}$= 7 | Time effect: F (23.00, 161.0) = 2.341 | **$p$ <0.0001** |
| | | | | - | - |
| | 300lux | Mixed-effects model (REML) | $n_{baseline}$= 10 | Interaction: F (23.00, 107.0) = 3.354 | **$p$ <0.0001** |
| | | | $n_{DLaN}$= 9 | Time effect: F (23.00, 207.0) = 12.56 | **$p$ <0.0001** |
| | | | | - | - |

"-": some tests can not be performed due to missing values around ZT13-16

| Table 6. Statistical difference in Figure 6 data | | | | | |
|---|---|---|---|---|---|
| **Description** | **Sexc** | **Light intensity** | **Statistical test** | **Number of pairs** | ***P* value/Exact *p* value** |
| Peak time of Waking | Male | 50 lux | Wilcoxon matched-pairs signed rank test | 10 | $p$=0.8809 |
| | | 100 lux | Wilcoxon matched-pairs signed rank test | 9 | $p$=0.8984 |
| | | 300 lux | Paired t test | 10 | $p$>0.9999 |
| | Female | 50 lux | Paired t test | 9 | **$p$=0.0062** |
| | | 100 lux | Paired t test | 7 | **$p$=0.035** |
| | | 300 lux | Paired t test | 9 | $p$=0.2623 |
| Peak time of NREM sleep | Male | 50 lux | Wilcoxon matched-pairs signed rank test | 10 | $p$=0.4023 |
| | | 100 lux | Paired t test | 9 | $p$=0.7811 |
| | | 300 lux | Wilcoxon matched-pairs signed rank test | 10 | $p$=0.2188 |
| | Female | 50 lux | Wilcoxon matched-pairs signed rank test | 9 | **$p$=0.0117** |
| | | 100 lux | Paired t test | 7 | **$p$=0.0212** |
| | | 300 lux | Paired t test | 9 | $p$=0.4379 |
| Peak time of REM sleep | Male | 50 lux | Wilcoxon matched-pairs signed rank test | 10 | $p$=0.5547 |
| | | 100 lux | Paired t test | 9 | $p$=0.7621 |
| | | 300 lux | Paired t test | 10 | $p$=0.1589 |
| | Female | 50 lux | Wilcoxon matched-pairs signed rank test | 9 | **$p$=0.0078** |
| | | 100 lux | Paired t test | 7 | $p$=0.0781 |
| | | 300 lux | Paired t test | 9 | $p$=0.1492 |
| Amplitude of Waking | Male | 50 lux | Wilcoxon matched-pairs signed rank test | 10 | **$p$=0.0039** |
| | | 100 lux | Paired t test | 9 | **$p$=0.0099** |
| | | 300 lux | Paired t test | 10 | **$p$=0.0167** |
| | Female | 50 lux | Paired t test | 9 | **$p$=0.0024** |
| | | 100 lux | Wilcoxon matched-pairs signed rank test | 7 | **$p$=0.0469** |
| | | 300 lux | Paired t test | 9 | $p$=0.2693 |
| Amplitude of NREM sleep | Male | 50 lux | Wilcoxon matched-pairs signed rank test | 10 | **$p$=0.0059** |
| | | 100 lux | Paired t test | 9 | **$p$=0.0342** |
| | | 300 lux | Paired t test | 10 | $p$=0.0562 |
| | Female | 50 lux | Paired t test | 9 | **$p$=0.001** |
| | | 100 lux | Paired t test | 7 | **$p$=0.0112** |
| | | 300 lux | Paired t test | 9 | $p$=0.143 |
| Amplitude of REM sleep | Male | 50 lux | Paired t test | 10 | **$p$=0.0458** |
| | | 100 lux | Paired t test | 9 | **$p$=0.0472** |
| | | 300 lux | Paired t test | 10 | $p$=0.5309 |
| | Female | 50 lux | Paired t test | 9 | **$p$=0.006** |
| | | 100 lux | Wilcoxon matched-pairs signed rank test | 7 | **$p$=0.0156** |
| | | 300 lux | Paired t test | 9 | $p$=0.09 |
